# Phenotypic Changes in Undifferentiated and Retinoic Acid Differentiated HL-60/S4 Cells with-or-without LBR Knockdown

**DOI:** 10.1101/2025.02.02.635278

**Authors:** Ada L. Olins, David Mark Welch, Igor Prudovsky, Donald E. Olins

## Abstract

Many functions of the nuclear envelope protein LBR (Lamin B Receptor) have been documented. LBR influences the shape of the nuclear envelope via its connections to underlying heterochromatin and to the lamina. LBR also functions as an essential enzyme in sterol biosynthesis. This study attempts to further define the LBR gene functions in a LBR knockdown cell line (HL-60/sh1), by employing Gene Set Enrichment Analysis (GSEA) to explore: Part 1) the effects of LBR knockdown on cell differentiation to granulocytes induced by retinoic acid (RA); Part 2) the effects of LBR knockdown on phenotypic differences between undifferentiated HL-60/sh1 and undifferentiated control cell lines (HL-60/S4 and HL-60/gfp). These GSEA studies are based upon previously published mRNA transcriptome data. In Part 1, the most significant effects were the increased <u>loss</u> of heterochromatin and the decreased histone methyltransferase activity in RA-differentiated HL-60/sh1 granulocytes, compared to HL-60/S4 and HL-60/gfp granulocytes. In addition, HL-60/sh1 ribosome structural protein transcripts were more <u>increased</u> during granulocyte differentiation, than observed in HL-60/S4 and HL-60/gfp. For many predicted phenotypes (e.g., senescence, migration, chemotaxis, phagocytosis and apoptosis), no significant differences were observed among the three granulocytic cell lines. In Part 2, comparisons were mainly between <u>undifferentiated</u> HL-60/sh1 and HL-60/S4 cells. We noted a significant increase in ribosomal protein and ribosomal RNA synthesis in sh1 0 versus to S4 0 cells. Of further interest, we observed that the position and number of nucleoli per cell appeared to differ between these two undifferentiated cell lines. In addition, GSEA results indicated a significant <u>gain</u> in heterochromatin and nucleosome formation in sh1 0 versus S4 0 cells. Microscopic imaging of undifferentiated sh1 0 cells compared to S4 0 cells indicated increased DAPI stained chromatin condensates surrounding the frequently central nucleoli, possibly reflecting the increased heterochromatin and the decreased LBR. Furthermore, the sh1 0 cell nuclei appeared “rounder” than the S4 0 cell nuclei. Evidence is presented supporting that the LINC Complex (“Linker of Nucleoskeleton and Cytoskeleton”) may play a role.

## Introduction

HL-60/S4 cells are an immortalized AML (Acute Myeloid Leukemia) cell line derived from the parent HL-60 cell line (Leung et al., 1992). They can be differentiated into granulocytes with retinoic acid (RA) and into macrophage with phorbol ester (TPA) following four days of drug exposure (Olins et al., 1998; Olins et al., 2009). Undifferentiated HL-60/S4 cells exhibit robust growth in suspension with a doubling time of ∼ 17 hours. RA-differentiated HL-60/S4 cells exhibit a gradual slowing of cell division and remain in suspension (Olins et al., 2001). Figure 1 illustrates that the initial “spheroid” nuclei becomes lobulated, with “ELCS” (i.e., nuclear <u>E</u>nvelope-<u>L</u>imited <u>C</u>hromatin <u>S</u>heets) bridging between the nuclear lobes (Olins et al., 1998). By four days of granulocyte differentiation, apoptosis becomes apparent (Olins et al., 1998). The protein composition of the granulocyte nuclear envelope also changes dramatically during differentiation, exhibiting an increase of Lamin B Receptor (LBR) and a decrease in Lamin A by day 4 (Mark Welch et al., 2017; Olins et al., 2001; Olins et al., 2009; Olins et al., 2010b).

**Figure 1.**
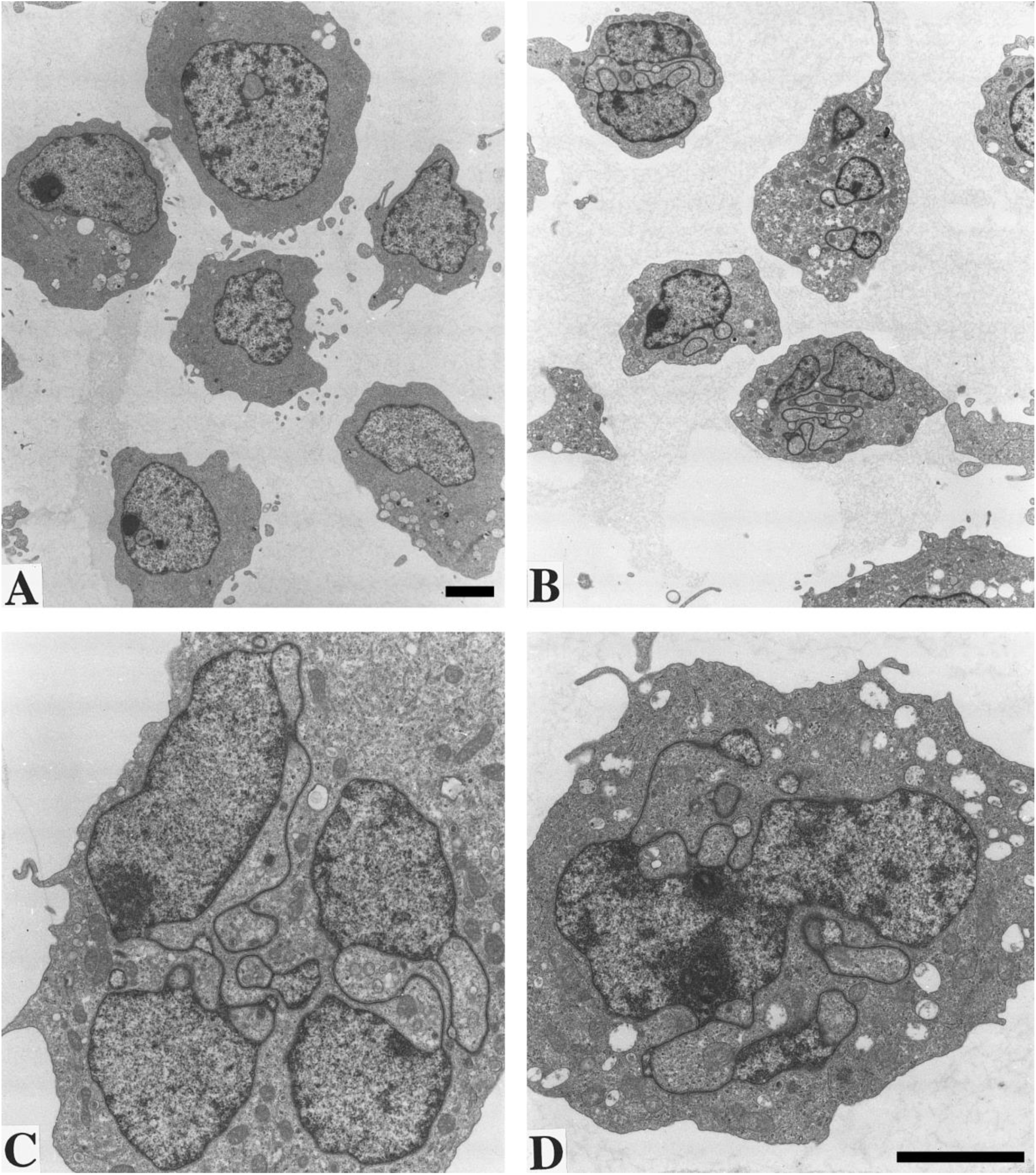
Electron micrographs of HL-60/S4, without RA treatment (A); after 4 days of RA induced differentiation (B, C, and D). Cells fixed in glutaraldehyde, OsO4, and stained with methanolic uranyl acetate and lead citrate. Nucleoli appear as strongly stained nuclear inclusions. ELCS, spanning between nuclear lobes, are best visualized in frames (C and D). Magnification Bars equal 3 µm. Image reproduced from (Olins et al., 1998)

The transcriptomes of HL-60/S4 undifferentiated (0), granulocytes (RA) and macrophage (TPA) have been a major tool for understanding the cellular phenotypes, since they were initially described (Mark Welch et al., 2017). Generally, the transcriptome data sets have been studied as Differential Gene Expression (DGE), being compared as Log_2_ Fold Changes (Log_2_FC) of normalized expression ratios; e.g., Log_2_FC (RA/0) or Log_2_FC (TPA/0) (Mark Welch et al., 2017). Most recently, we have examined the DGE changes between an “unmodified” HL-60 cell line (HL-60/S4), a stable lentiviral LBR knockdown subline of HL-60 (HL-60/sh1), and a stable cell line containing a gfp-lentivirus lacking the LBR hairpin (HL-60/gfp) as an additional control (Olins et al., 2010a) (Table 1). This data was employed to identify enriched Gene Ontology (GO) terms, in order to identify phenotypic changes during differentiation in RA-treated LBR “knockdown” HL-60/sh1 cells (Mark Welch et al., 2024).

**Table 1.** Transcription level changes in HL-60/S4, HL-60/sh1 and HL-60/gfp cell lines after differentiation with RA. Cell lines: S4 cells are the “unmodified” control cell line; sh1 cells are a “lentivirus-LBR short hairpin knockdown” cell line; gfp cells are a “gfp-lentivirus-only” control cell line (Olins et al., 2010a). DGE: Differential Gene Expression categories. Total: Total number of <u>significant</u> genes (PPDE≥ 0.95). # of genes: number of genes in each category of DGE (Increased, Decreased or Unchanged). % of Total: percentage of each category of DGE in the total number of significant genes.

| RA/0 |  |  |  |  |  |  |
| --- | --- | --- | --- | --- | --- | --- |
| Title | S4 |  | sh1 |  | gfp |  |
| DGE | # of genes | % of total | # of genes | % of total | # of genes | % of total |
| Increased | 3,989 | 23.7 | 4,134 | 24.6 | 4,469 | 26.6 |
| Decreased | 3,514 | 20.9 | 4,030 | 24 | 4,582 | 27.2 |
| Unchanged | 9,319 | 55.4 | 8,658 | 51.5 | 7,771 | 46.2 |
| Total | 16,822 |  | 16,822 |  | 16,822 |  |

Part 1 of the present study is focused upon the RA differentiated cellular phenotypic changes related to relative changes of LBR transcript levels in the HL-60/S4, sh1 and gfp granulocytes. These studies are sequential to an earlier publication that describes the microscopic changes of nuclei during RA-induced differentiation of LBR knockdown (HL-60/sh1) cells (Olins et al., 2010a).

The most obvious consequence of LBR knockdown in HL-60/sh1 cells is the <u>absence</u> of nuclear lobulation and ELCS formation in the RA-differentiated cells (Figure 2). This absence of nuclear lobulation is reminiscent of neutrophil nuclei observed in the blood of humans with Pelger-Huet anomaly (Hoffmann et al., 2002; Olins et al., 2010a; Turner and Schlieker, 2016; Young et al., 2021). Two additional phenotypic changes have been observed in RA-treated HL-60/sh1 cells (Mark Welch et al., 2024): 1) Golgi associated vesicles immunostained with anti-TRIP11 are less prevalent in sh1 cells, than in control RA-treated HL-60/S4 cells; 2) Interphase nucleoli in both untreated and RA-treated sh1 cells frequently appear to be <u>single centrally-located</u> nuclear organelles; whereas, in control untreated and control RA-treated HL-60/S4 cells, interphase nuclei frequently exhibit multiple nucleoli, often adjacent to the nuclear envelope.

**Figure 2.**
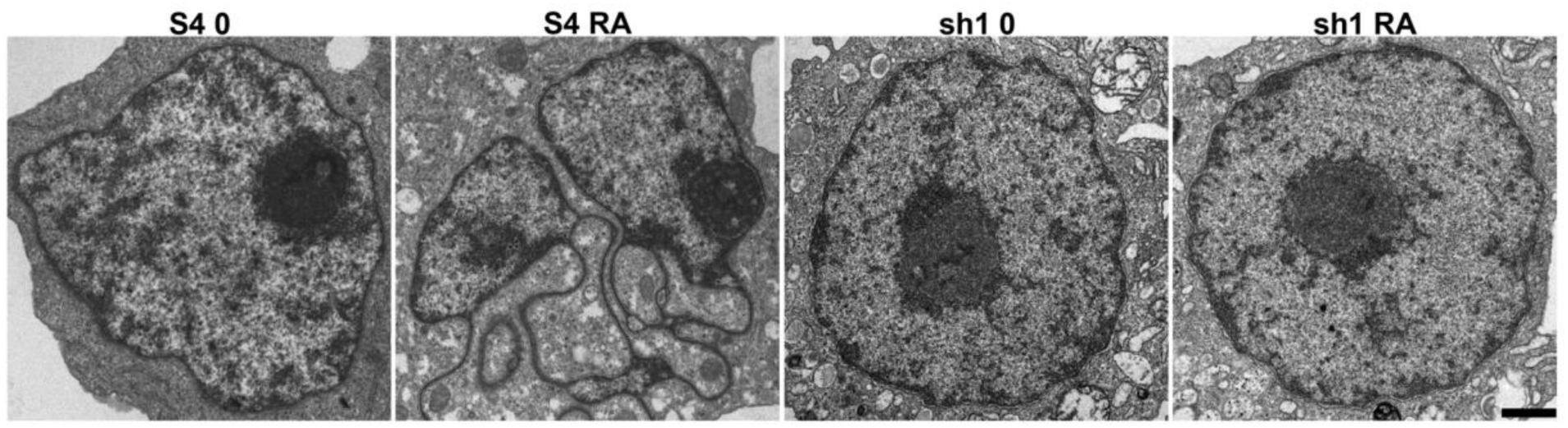
Thin-section electron microscopy of undifferentiated (0) and RA-differentiated S4 and sh1 cells. Note the appearance of a lobulated nucleus and chromatin sheets (ELCS) in between the nuclear lobes in the RA-treated S4 cells (S4 RA) and their absence in the RA-treated sh1 cells (sh1 RA). Note the positions of the densely stained nucleoli adjacent to the NE in the S4 cells, and in the center of the nucleus in sh1 cells. Magnification bar: 1 μm. These images have been previously published (Olins et al., 2010a).

The present study represents an extension of these prior publications, utilizing Gene Set Enrichment Analysis (GSEA) as a method to predict RA-differentiated phenotypic functional changes resulting from LBR knockdown in HL-60/sh1 cells, compared to the similarly treated S4 and gfp cell lines. GSEA is a computational method that determines whether a *pre-defined* set of genes is significantly over-represented (enriched) among all differentially expressed genes between two biological states or “phenotypes” (Mootha et al., 2003; Subramanian et al., 2005). For example, the “phenotypes” RA-treated HL-60/S4 versus untreated (0) HL-60/S4 cells. We have previously employed GSEA to describe the multifaceted phenotypes of TPA (phorbol ester)-treated versus untreated (0) HL-60/S4 cells (Olins et al., 2024). This earlier analysis substantiated that HL-60/S4 macrophages are very likely to be in senescence, much more so than for undifferentiated HL-60/S4 cells or for RA-differentiated HL-60/S4 granulocytes.

## Materials and Methods

### Cell Cultivation

HL-60/S4 cells can be purchased from ATCC (CRL-3306). They are cultivated in RPMI 1640 medium + 10% (unheated) Fetal calf serum + 1% Pen/Strep/Glut. We employ T-25 (6 ml) or T-75 (18 ml) flasks, lying horizontal to maximize surface area. The flasks are incubated humidified at 37° C with 5% CO2. HL-60/sh1 and HL-60/gfp were generated and cultivated as described (Olins et al., 2010a). Unfortunately, the HL-60/gfp culture was lost in a laboratory accident. However, this cell loss occurred after undifferentiated and differentiated mRNA transcriptomes were generated. The HL-60/sh1 and HL-60/gfp cell lines were cultivated in normal growth medium plus 1µg/ml of puromycin, to maintain selection pressure on the lentivirus infected cell lines. Puromycin was not present during cell differentiation.

### Granulocyte Differentiation

Stock solution: 5 mM retinoic acid (RA) in 100% ethanol. We use Sigma-Aldrich R2625 (MW=300.4). Store the RA stock solution wrapped in aluminum foil at -20° C. In a T-25 flask containing 5 ml of HL-60/S4 diluted with fresh medium to ∼2x10^5^ cells/ml, add 1μl of 5 mM RA. Wrap the flask with aluminum foil before placing in the incubator, to protect against light exposure. The cells will continue to replicate (more slowly) during differentiation, gradually stopping. “Crawling” granulocytes begin to appear by day 3. After day 4, apoptosis becomes apparent. The cells never attach strongly to the flask bottom.

### Gene Set Enrichment Analysis (GSEA)

For a detailed description of this method as applied to HL-60/S4, see Materials and Methods (Olins et al., 2024).

### RNA purification, RNA-Seq, Data analyses and presentation, Microscopy

These topics have all been described in an earlier Materials and Methods (Mark Welch et al., 2024). This cited BioRxiv preprint contains two Supplementary Tables that were also employed in the present article: 1) “Table S1 3-way.xlxs”, which was employed to compare the relative transcript levels of specific genes of the three cell lines (i.e., S4, sh1and gfp) in their undifferentiated <u>or</u> RA-differentiated cell states. 2) “Table S2 pairwise.xlxs”, which was utilized to compare the relative transcript levels of specific genes in their RA-treated versus untreated (RA/0) states, within a particular cell line.

### Antibodies

The only antibodies employed within this study were: 1) Active Motif (Catalogue #39239) Rabbit anti-H3K9me2; Active Motif (Catalogue #39161) Rabbit anti-H3K9me3. Both antibodies were employed for immunofluorescent staining and immunoblotting in Figures 8 and 9. As recommended by Active Motif, the same dilution factor (1:1000) was employed for both antibodies in all experiments.

## Results

The first part of the Results section is primarily a study of the functional consequences to RA-induced cell differentiation in the three cell lines (HL-60/S4, HL-60/sh1 and HL-60/gfp). The goal is to understand how LBR knockdown has affected the differentiation process. The second part of the Results section is primarily a comparative study of the three undifferentiated cell lines (HL-60/S4, HL-60/sh1 and HL-60/gfp). The goal of this study is to understand how LBR knockdown has affected the structure and function of the undifferentiated cell lines.

### Part One

#### Cellular Senescence appears to be similarly reduced in the three cell line Granulocytes

In a previous study we reported that TPA-treated HL-60/S4 macrophages exhibited distinct characteristics of senescent cells (Olins et al., 2024). In the present study, we compare selected genes in RA-differentiated cell lines HL-60/S4, HL-60/sh1 and HL-60/gfp with the same genes from TPA-treated HL-60/S4 macrophages (Table 2). When examining four genes involved in the cell cycle, it is clear that transcript levels in the RA-treated cells (granulocytes), with-or-without LBR knockdown, resemble each other, more than they resemble the TPA-treated cells (senescent macrophages). These results are consistent with the almost immediate cessation of cell division in the TPA-treated HL-60/S4 macrophages, and the slower cessation of cell division in RA-treated granulocytes (Mark Welch et al., 2017).

**Table 2.** Log_2_ Fold Changes of selected gene transcripts from the mRNA transcriptomes of RA-treated HL-60/S4, HL-60/sh1 and HL-60/gfp and TPA-treated HL-60/S4, each compared to the appropriate untreated (0) cells (Mark Welch et al., 2017; Mark Welch et al., 2024).

| STABLE CESSATION OF THE CELL CYCLE |  |  |  |  |  |  |
| --- | --- | --- | --- | --- | --- | --- |
|  |  |  | Log<sub>2</sub>FC |  |  |  |
| Gene Name | Code | Inhibition of | S4 RA/0 | sh1 RA/0 | gfp RA/0 | S4 TPA/0 |
| p21 | CDKN1A | CDK2&4 | 3.78 | 3.16 | 3.63 | 10.68 |
| p16 | CDKN2A | CDK4 | -0.61 | 0.09 | -0.37 | 1.6 |
| Telomerase | TERT | -- | -5.49 | -4.12 | -5.19 | -10.06 |
| Cyclin A2 | CCNA2 | -- | -0.58 | -0.41 | -0.56 | -3.46 |

GSEA enrichment plots of the four differentiated cell States described in Table 2, tested against the SenMayo senescence gene set (125 genes) (Saul et al., 2022) are shown in Figure 3. All the plots have “p-nominal” scores of 0.0, indicating that the NES values are statistically significant. The higher NES (Normalized Enrichment Score) for TPA-differentiated HL-60/S4 macrophages supports the contention that enrichment of transcripts related to senescence is greater in this cell line, than in RA-differentiated granulocytes with-or-without LBR knockdown.

**Figure 3.**
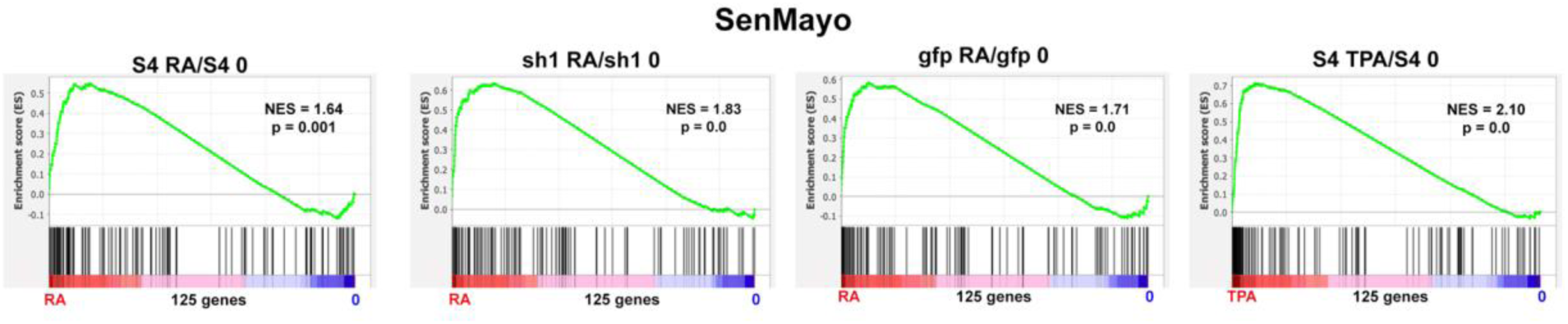
GSEA enrichment plots of the SenMayo gene set (125 genes). Each analysis compares DGE of the same cell line (i.e. S4, sh1, gfp) under two different phenotype conditions (i.e., RA vs 0). For each plot the two conditions represent the boundaries of the ranked genes.

#### Numerous Granulocyte Phenotypes exhibit similar GSEA enrichment in the three RA-treated HL-60 cell lines

Given the drastic changes in nuclear morphology observed in the RA-treated HL-60 cell lines (Figures 1 and 2), we decided to explore whether there are phenotypic differences in the functions of HL-60/S4, HL-60/sh1 and HL-60/gfp, including cell differentiation, activation, chemotaxis, migration and phagocytosis. MSigDB gene sets are available for these cellular functions. Using these chosen gene sets, there are only minor transcriptome differences (NES and p-nominal values) comparing the three RA-treated cell lines (Figure 4 and Table 3). These GSEA analyses constitute <u>predictions</u> of cellular functions, since they are based upon relative levels (RA/0) of relevant gene transcripts. Future studies can search for differences in chemotaxis and phagocytosis in the live differentiated granulocytes.

**Figure 4.**
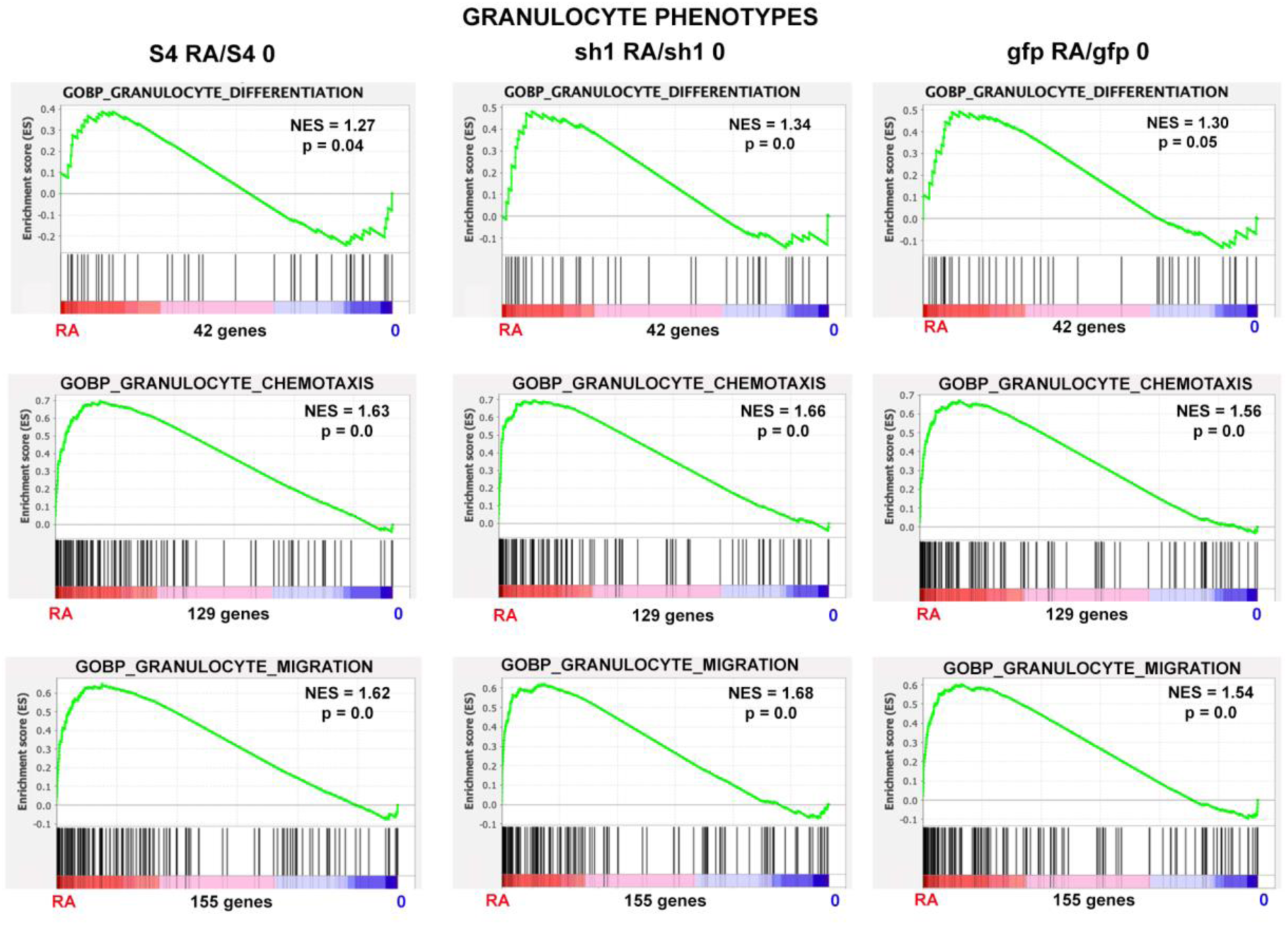
GSEA enrichment plots of three MSigDB gene sets (rows) for the three different cell lines shown in columns (HL-60/S4, HL-60/sh1 and HL-60/gfp), with-or-without RA-induced differentiation. Analysis of additional MSigDB gene sets revealed no significant differences between the three cell lines (Table 3; GSEA enrichment plots are not shown for all gene sets). In other words, by these Granulocyte Phenotype criteria, the three cell lines (HL-60/S4, sh1 and gfp) are remarkably similar, independent of the LBR knockdown.

**Table 3.**
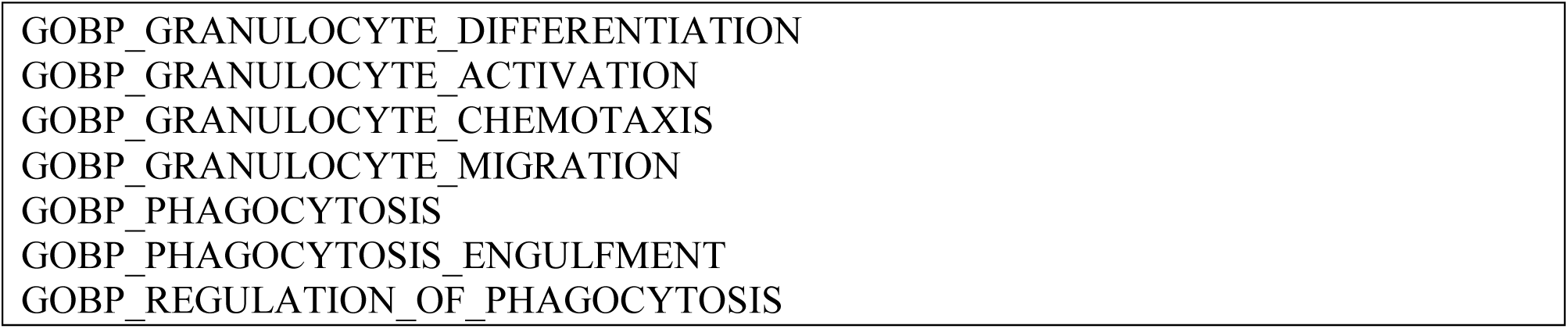
MSigDB gene sets that show no significant Granulocyte Phenotypic differences among RA-treated HL-60/S4, HL-60/sh1 and HL-60/gfp.

#### The Apoptosis Phenotype displays similar GSEA enrichment in all three RA-treated HL-60 cell line Granulocytes

RA treatment of HL-60/S4 cells result in obvious apoptosis by day 4 of exposure (Olins et al., 1998). During induced differentiation of HL-60/S4 cells to granulocyte form, transcript levels of LBR increases ∼3 fold (Mark Welch et al., 2024; Olins et al., 1998). An important question is whether LBR knockdown in HL-60/sh1 cells treated with RA results in a reduction of the apoptosis phenotype. Figure 5 compares GSEA enrichment plots of apoptosis genes in the three RA-treated cell lines HL-60/S4, HL-60/sh1 and HL-60/gfp. The NES values for these cell lines are very similar with significant p values. It is clear that the considerable reduction of LBR in HL-60/sh1 granulocytes (Mark Welch et al., 2024) does <u>not</u> reduce the enrichment of the apoptosis phenotype.

**Figure 5.**
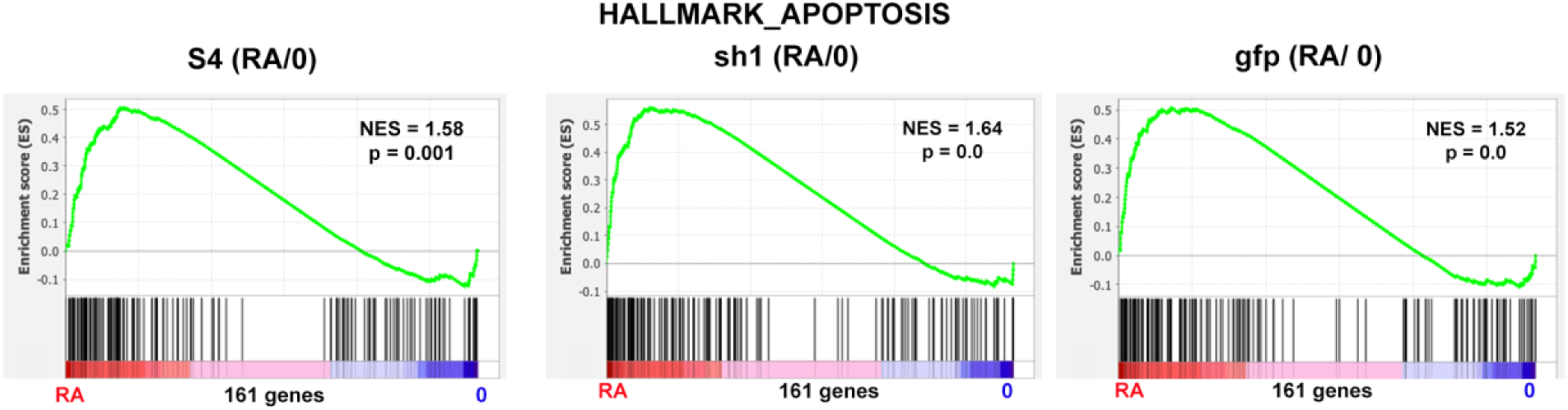
GSEA enrichment plots of the MSigDB gene set “HALLMARK_APOPTOSIS” for each cell line with-or-without RA.

#### HL-60/sh1 Granulocytes display a greater <u>loss</u> of Heterochromatin, compared to the S4 and gfp Granulocytes

In our previous publication on the GSEA phenotypic properties of TPA-differentiated HL-60/S4 macrophages (Olins et al., 2024), we observed that these macrophages exhibited <u>reduced</u> (i.e., not enriched) levels of heterochromatin-related transcripts, compared to the undifferentiated (0) HL-60/S4 cells. This observation was consistent with earlier publications from other laboratories, which presented evidence that senescent cells exhibit a reduction in heterochromatin (Parry et al., 2018; Rocha et al., 2022; Sławińska and Krupa, 2021; Swanson et al., 2015; Zhang et al., 2021). This earlier finding provoked us to examine whether RA-differentiated HL-60 granulocytes also show a reduction in heterochromatin, and whether there are differences among the different HL-60 cell lines (e.g., S4, sh1 and gfp); see Figure 6. Table 4 presents calculations from GSEA enrichment plots for three types of heterochromatin: e.g., (Global) Heterochromatin (shown in Figure 6); Pericentric (Constitutive) Heterochromatin; Facultative Heterochromatin. In all cases of RA-differentiated granulocytes, the NES value for RA vs 0 phenotypes were <u>more reduced</u> in HL-60/sh1 cells, than in HL-60/S4 and HL-60/gfp cells. This observation argues that heterochromatin is decreased during granulocytic differentiation, with the greatest loss in HL-60/sh1 cells.

**Figure 6.**
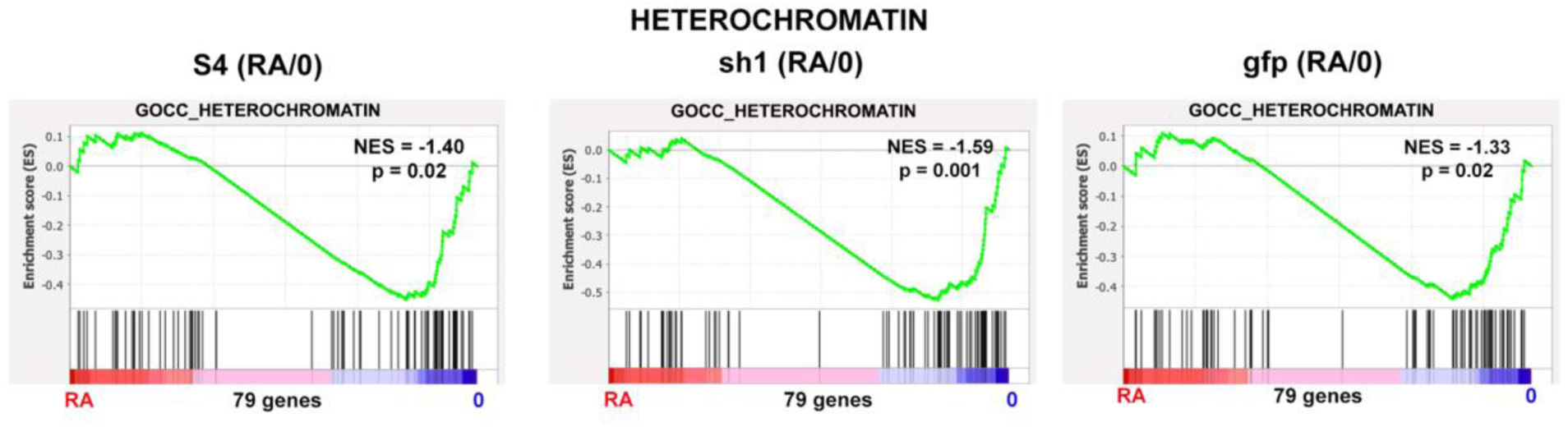
GSEA enrichment plots of the MSigDB gene set GOCC_HETEROCHROMATIN for the three cell lines (HL-60/S4, HL-60/sh1 and HL-60/gfp), for each cell line with-or-without RA. These results revealed that all three cell lines exhibited decreased global heterochromatin (i.e., negative NES), with the largest decrease in HL-60/sh1 granulocytes (Table 4).

**Table 4.** GSEA results for three types of heterochromatin. For all three types of heterochromatin, sh1 RA/0 exhibits a greater decrease in NES values than S4 RA/0 or gfp RA/0.

| CELLS | NES | Nominal p |
| --- | --- | --- |
| S4 RA/0 | -1.40 | 0.02 |
| sh1 RA/0 | -1.59 | 0.001 |
| gfp RA/0 | -1.33 | 0.02 |

GOCC\_PERICENTRIC\_HETEROCHROMATIN. 30 genes
| CELLS | NES | Nominal p |
| --- | --- | --- |
| S4 RA/0 | -1.41 | 0.04 |
| sh1 RA/0 | -1.65 | 0.01 |
| gfp RA/0 | -1.36 | 0.08 |

| CELLS | NES | Nominal p |
| --- | --- | --- |
| S4 RA/0 | -1.19 | 0.17 |
| sh1 RA/0 | -1.62 | 0.003 |
| gfp RA/0 | -1.28 | 0.10 |

In order to investigate the basis of this “sh1-specific” effect on HL-60 granulocytes, we displayed the GOCC_HETEROCHROMATIN Leading Edges of the S4, sh1 and gfp cell lines (Table S1). Leading Edge analysis identifies those genes within a specific GSEA gene set that contribute most to the enrichment score (ES). In this case, comparing the three different cell lines’ Leading Edges from GOCC_HETEROCHROMATIN, we can speculate which genes are candidates for the sh1-specific effect. Shown in alphabetic order, Table S1 identifies 8 candidate genes (i.e., BAZ1A, BAZ1B, CBX1, CENPC, INCENP, MACROH2A1, MPHOSPH8 and ZNF618) that are present in the sh1 Leading Edge, but absent from the S4 and gfp Leading Edges. These 8 genes <u>all</u> appear to be transcriptionally downregulated in the RA-differentiated HL-60/sh1 granulocytes (Figure 7). Of particular interest is CBX1, which can interact within the N-terminal peptide region of LBR at the “HP1 binding motif” (Olins et al., 2010b). The reduction of LBR in HL-60/sh1 cells (Mark Welch et al., 2024), combined with a reduction of CBX1 may “weaken” the nuclear envelope-heterochromatin “bridge”. The other 7 candidate genes are involved in chromatin remodeling, binding to centromeric regions and promoting repression of transcription. Their reduced transcription might also contribute to dismantling heterochromatin. Furthermore, the combined decrease in CBX1 and CENPC in HL-60/sh1 cells may indicate that pericentric heterochromatin has one particular target (i.e., kinetochore) with a disabled function.

**Figure 7.**
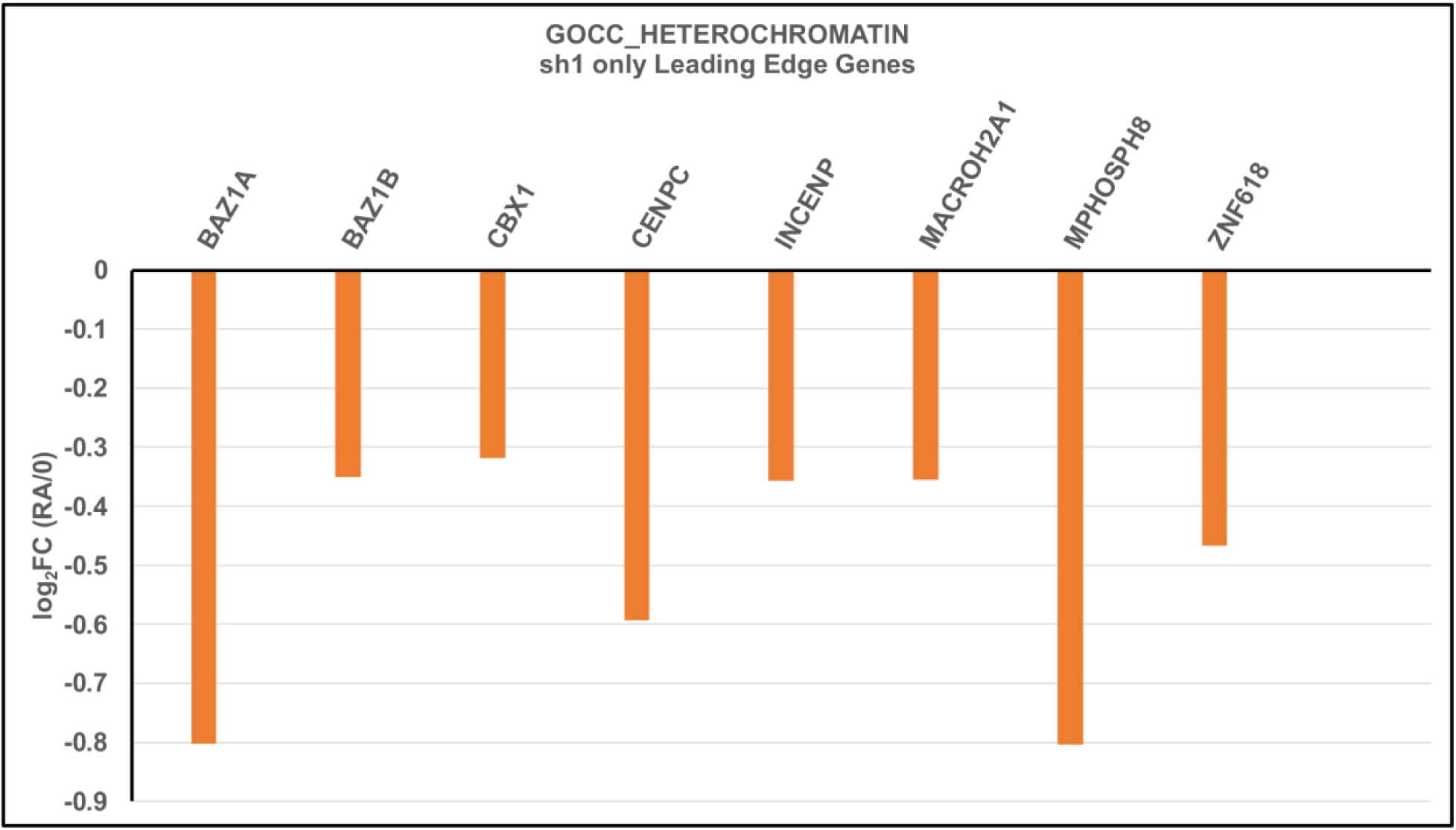
Downregulation (reduced transcript levels) of specific genes identified within the Leading Edge of GOCC_HETEROCHROMATIN for HL-60/sh1 granulocytes, but absent from the Leading Edge of HL-60/S4 and HL-60/gfp granulocytes (Table S1).

Table S1 also displays the Leading Edges of two other heterochromatin MSigDB gene sets listed in Table 4: GOCC_PERICENTRIC_HETEROCHROMATIN and GOBP_FACULTATIVE_HETEROCHROMATIN_FORMATION. The gene codes highlighted in red represent the genes exclusively downregulated in sh1 cells.

PERICENTRIC_HETEROCHROMATIN shares 4 candidate genes (i.e., BAZ1B, CBX1, CENPC and ZNF618) present in the sh1 Leading Edge and absent from the S4 and gfp Leading Edges. FACULTATIVE HETEROCHROMATIN shares 3 genes (MACROH2A1, MPHOSPH8 and SMARCA5). These latter three genes are (respectively) associated with transcription repression, recognition of H3K9 methylation and nucleosome remodeling (www.genecards.org). Overall, the transcriptome analyses are consistent with decreased heterochromatin in the RA-treated S4, sh1 and gfp cell lines, especially apparent in the HL-60/sh1 cell lines.

#### HL-60/S4, HL-60/gfp and, especially, HL-60/sh1 Granulocytes exhibit decreased levels of Histone Methylation enzymes

Among the factors that regulate the existence of heterochromatin is epigenetic modification of nucleosomes, which includes histone methylation, acetylation and phosphorylation. Histone methylation is a complex field of study, which has been recently comprehensively reviewed (Fukuda et al., 2023; Padeken et al., 2022). Generally, histone lysine methylation (e.g., H3K9me1, 2 and 3) correlates with transcription repression (downregulation) in heterochromatin regions along chromosomes. However, some histone methylation (e.g., H3K4me1) has been implicated in upregulation of transcription (Wang and Ren, 2024). Epigenetic changes in chromatin are accomplished by a plethora of nucleosome modifying enzymes. Only a few examples are discussed in the present study.

Among the best studied examples of mammalian histone lysine methylation of H3K9me, present within pericentric heterochromatin, results from the expression of the SUV39H1 and SUV39H2 genes, which transcribe the methyltransferase enzymes responsible for formation of H3K9me2 and 3. These methylated histone lysines bind to the heterochromatin proteins CBX1, 2 and 3. Figure 8 presents images of S4 0, S4 RA, sh1 0 and sh1 RA cells, immunostained with rabbit anti-H3K9me2 and anti-H3K9me3. Figure 9 presents immunoblot images of SDS extracts from the same set of cells, employing the same rabbit anti-H3K9me2 and anti-H3K9me3 antibodies as used in Figure 8. The extract loads for the SDS-PAGE immunoblot were identical to those employed earlier, see Figure 3 of (Mark Welch et al., 2024).

**Figure 8.**
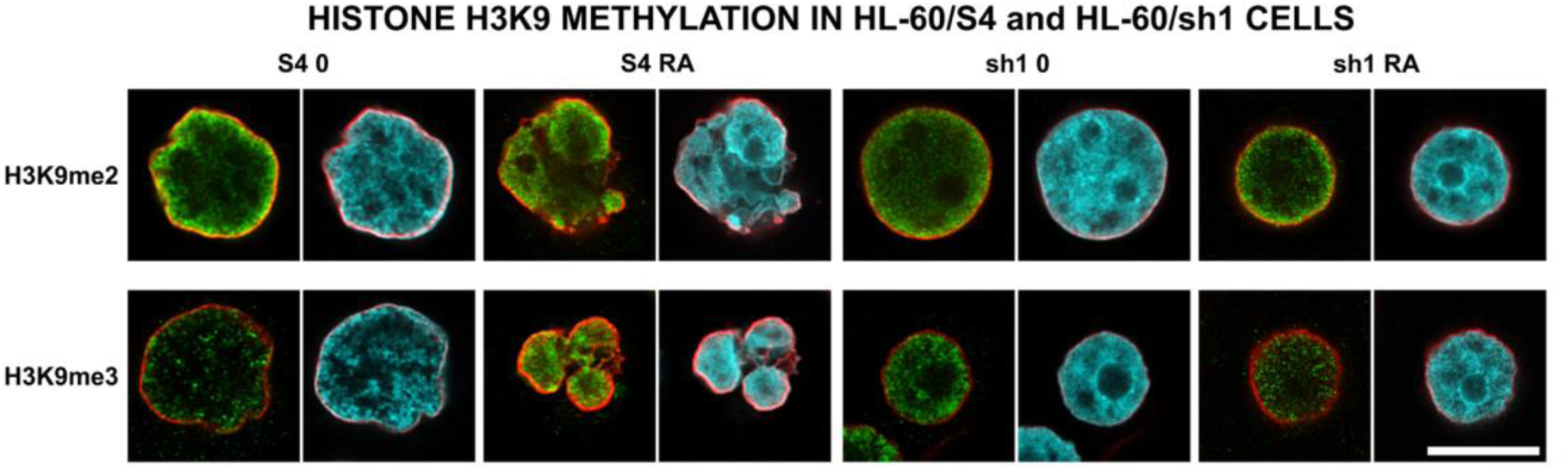
Images of S4 0, S4 RA, sh1 0 and sh1 RA cells, immunostained with rabbit anti-H3K9me2 and anti-H3K9me3. Color scheme: Red, epichromatin (Olins and Olins, 2018); Green, histone methylation antibody staining; Cyan, DAPI staining. Because of image deconvolution, the relative staining intensities of each antibody, comparing different cell types, are not preserved. This judgement should be based upon the immunoblotting results (Figure 9). The value of the fluorescent images is to show the nuclear locations where the H3K9 methylation epitopes are exposed. It appears that H3K9me2 is more localized near the nuclear envelope; H3K9me3 appears to be more uniformly spread within the interphase nucleus. Magnification bar: 10 μm.

**Figure 9.**
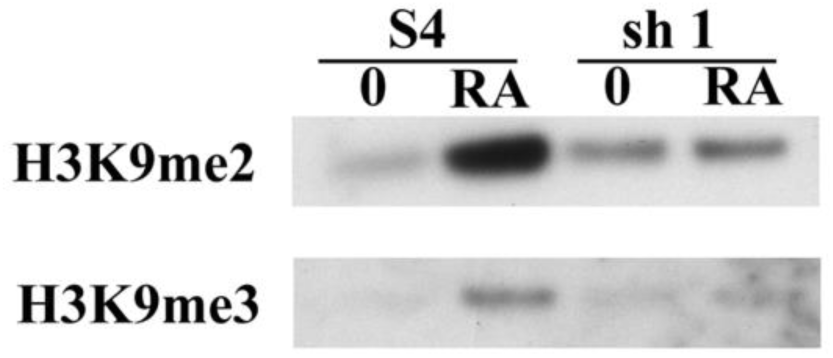
Immunoblot images of SDS extracts from S4 0, S4 RA, sh1 0 and sh1 RA cells, reacted with rabbit anti-H3K9me2 and anti-H3K9me3 antibodies. Note the clear reduction of H3K9me2 and H3K9me3 modifications in the sh1 cell extracts, compared to the RA-treated HL-60/S4 extracts. The extract loads for the SDS-PAGE immunoblot were identical to those employed earlier; see Figure 3 of (Mark Welch et al., 2024), which shows a Coomassie Blue stained gel with the same extract loads adjacent to molecular weight (MW) markers (kDa).

MSigDB gene sets are available for evaluating the enrichment of histone lysine methylation enzymes within appropriate transcriptomes. Figure 10 presents an example of plots for the MSigDB gene set (GOMF_HISTONE_METHYLTRANSFERASE_ACTIVITY), revealing the significant downregulation of histone methylation enzyme transcripts in the three cell lines HL-60/S4, HL-60/sh1 and HL-60/gfp, following RA-induced differentiation to granulocytes. Table 5 presents the GSEA enrichment results for four different GSEA gene sets of histone methylation enzymes. For all the different gene sets, sh1 RA/0 exhibits the greatest <u>decrease</u> in NES values, compared to S4 RA/0 and to gfp RA/0.

**Figure 10.**
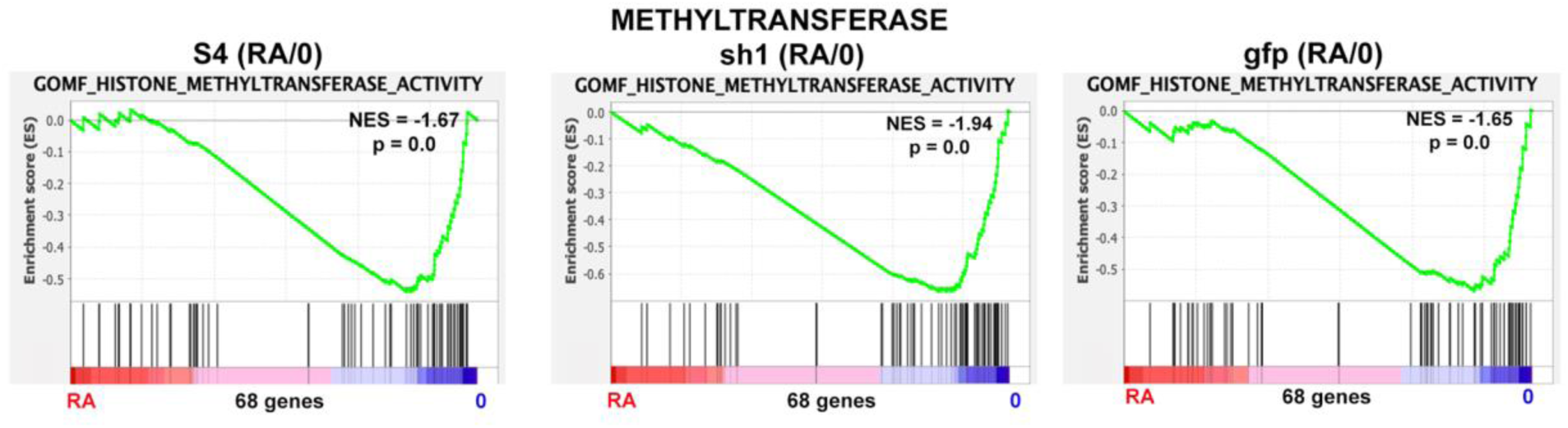
GSEA enrichment plots of the MSigDB gene set GOMF_HISTONE_METHYLTRANSFERASE_ACTIVITY for the three cell lines (HL-60/S4, HL-60/sh1 and HL-60/gfp) with-or-without RA. These results revealed that all three cell lines exhibited decreased histone methylation transcriptomes (i.e., negative NES), with the largest decrease in HL-60/sh1 granulocytes (Table 5).

**Table 5.**
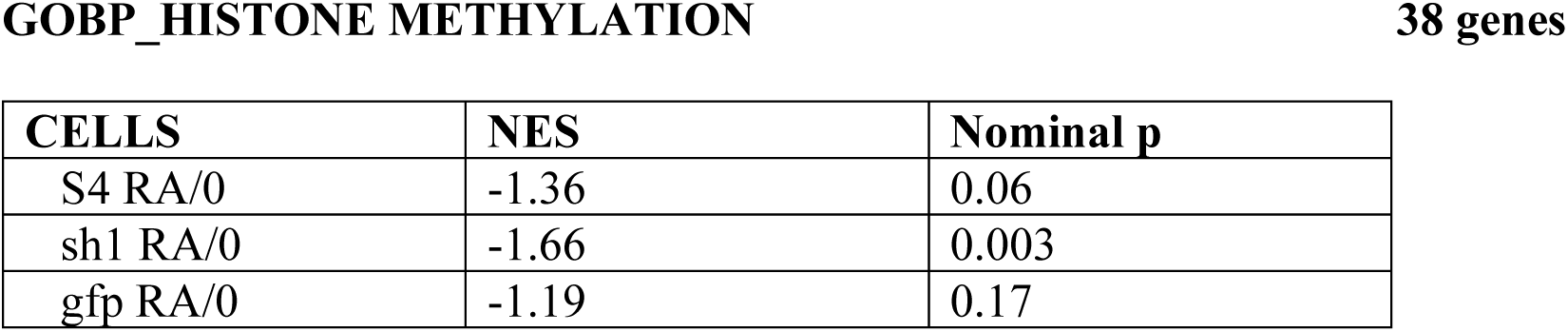
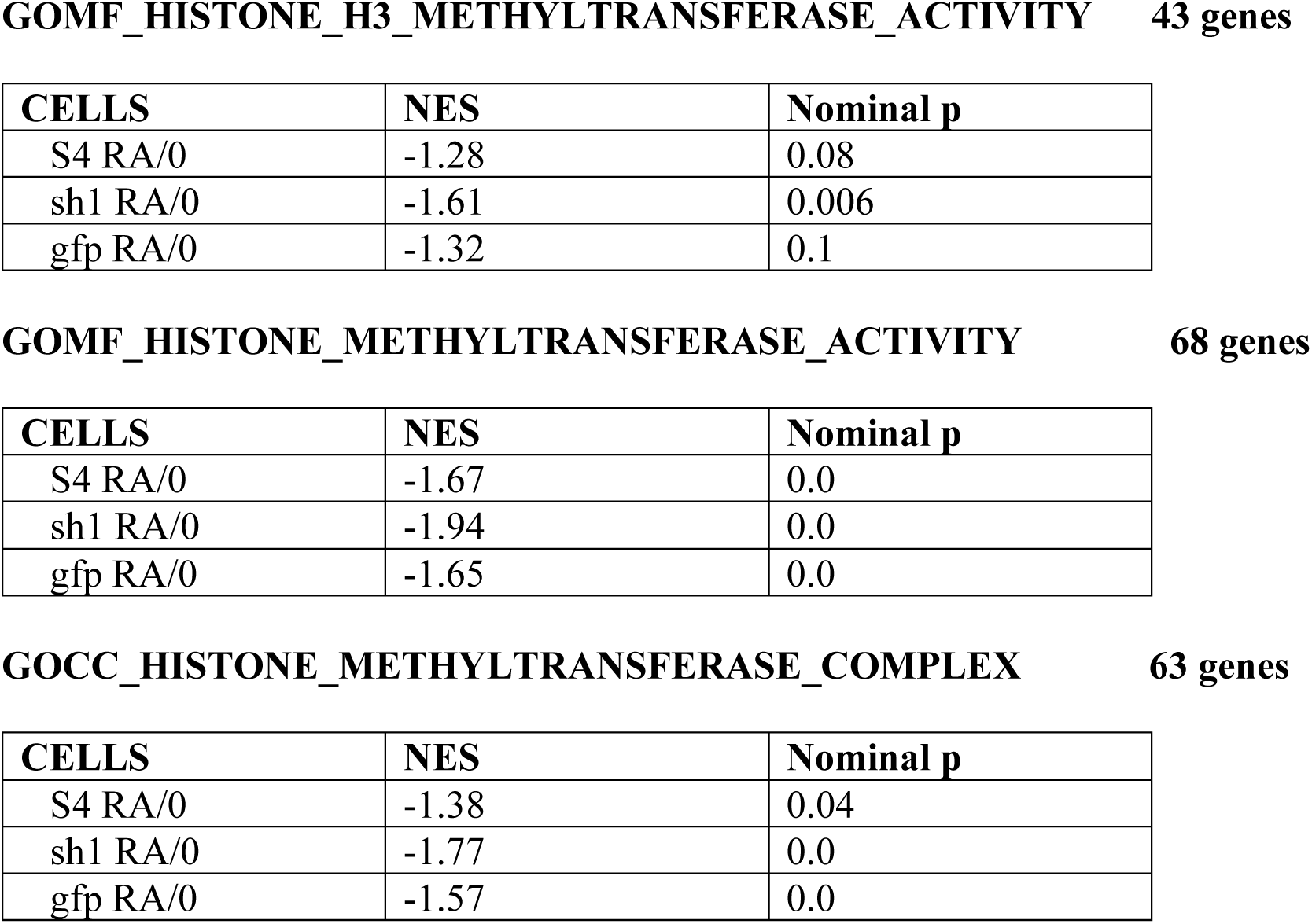
GSEA results for four different gene sets of histone methylation enzymes. For all four different gene sets, sh1 RA/0 exhibits the greatest decrease in NES values, compared to S4 RA/0 and to gfp RA/0.

The Leading Edges of these four different MSigDB gene sets were compared, looking for genes in sh1 cells that might account for their greater “loss” of histone methylation (Table S2). The SUV39H1 and SUV39H2 genes are clearly downregulated in two of the Leading Edges (GOMF_HISTONE_H3_METHYLTRANSFERASE_ACTIVITY and GOMF_HISTONE_METHYLTRANSFERASE_ACTIVITY), but not in the other two, implying the involvement of other lysine methyltransferases. Of interest, comparing the Leading Edges of the four different MSigDB gene sets (Table S2) reveals <u>only one</u> common downregulated candidate gene: KMT2A. Of possible interest, various chromosome translocations of this histone methyltransferase gene have been implicated in causing some Acute Myeloid Leukemia (AML). See: www.genecards.org, KMT2A

#### Based upon GSEA, HL-60/sh1 granulocytes exhibit significantly greater enrichment of ribosomal proteins, compared to HL-60/S4 and HL-60/gfp granulocytes

In our earlier study (Mark Welch et al., 2024), employing over-representation analysis of GO terms, “Table 2/GO Terms <u>Only-sh1-up</u>”, the most statistically significant GO Term was “Ribosome”. There are 19 “<u>Only-sh1-up”</u> genes that overlapped with the “Ribosome” gene set; the vast majority were for ribosome structural proteins (Table S3). However, in the present study, employing GSEA enrichment (Table 6), we have encountered an apparent paradox that questions whether there are increased amounts of functional ribosomes. GSEA gene sets GOBP_RIBOSOME_ASSEMBLY and GOBP_RIBOSOME_BIOGENESIS indicate downregulation of ribosomes in S4, sh1 and gfp cells during differentiation with RA, although HL-60/sh1 cells show the <u>least</u> downregulation.

**Table 6.** GSEA enrichment plots parameters for two different gene sets of ribosome formation. For the two different gene sets, sh1 RA/0 exhibits the <u>least decrease</u> in NES values, compared to S4 RA/0 and to gfp RA/0.

| CELLS | NES | Nominal p |
| --- | --- | --- |
| S4 RA/0 | -2.10 | 0.0 |
| sh1 RA/0 | -1.51 | 0.02 |
| gfp RA/0 | -1.87 | 0.0 |

| CELLS | NES | Nominal p |
| --- | --- | --- |
| S4 RA/0 | -2.63 | 0.0 |
| sh1 RA/0 | -2.31 | 0.0 |
| gfp RA/0 | -2.57 | 0.0 |

To analyze more deeply the basis for this observation, we interrogated the GSEA gene sets: GOCC_RIBOSOME and KEGG_RIBOSOME, comparing the HL-60/sh1 RA versus 0 phenotypes (Figure 11, Top Row). These GOCC and KEGG gene set Leading Edges are highly enriched for the gene codes of ribosomal structural proteins. In contrast, the Leading Edges of GOBP_RIBOSOME_ASSEMBLY and GOBP_RIBOSOME_BIOGENESIS exhibit a paucity of ribosomal structural genes and an enrichment in non-ribosomal structural genes (Table S4).

**Figure 11.**
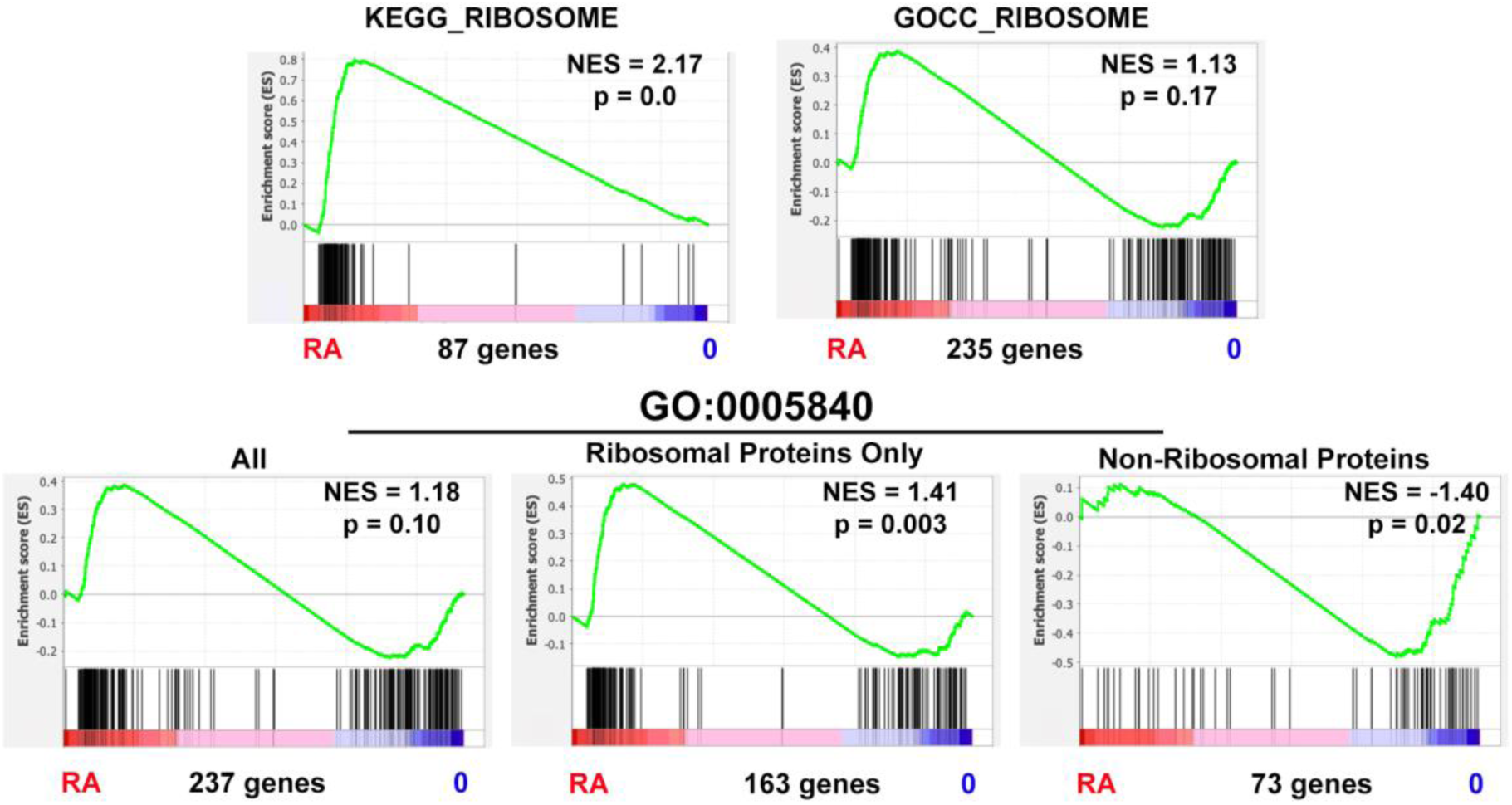
GSEA enrichment plots comparing the HL-60/sh1 RA versus 0 phenotypes. Top Row: GOCC_RIBOSOME and KEGG_RIBOSOME. Bottom Row: gene sets based upon division of the GO term GO:0005840 “Ribosome” into three groups: “ribosomal protein only” (“rp”, 163 genes); “non-ribosomal protein” (“nrp”, 73 genes); and their composite group “ALL” (237 genes).

Finally, to evaluate the role of these gene categories, we divided the genes within the GO term GO:0005840 “Ribosome” into two groups: “ribosomal protein only” (“rp”, 163 genes); “non-ribosomal protein” (“nrp”, 73 genes). These two gene groups and their composite group “ALL” (237 genes) were employed as the gene sets for additional GSEA enrichment plots, comparing the HL-60/sh1 RA versus 0 phenotypes (Figure 11, Bottom Row). Table 7 presents GSEA plot parameters for the five different gene sets: KEGG; GOCC; ALL; ribosomal protein only (rp); non-ribosomal protein (nrp). These results were derived from analysis of the transcriptomes (RA/0) of HL-60/S4, HL-60/sh1 and HL-60/gfp granulocytes. Our conclusion from all of these comparisons is that RA-induced differentiation results in increased transcripts for ribosomal structural proteins in HL-60/sh1 cells, compared to decreases in HL-60/S4 and HL-60/gfp cells. This conclusion is also consistent with an earlier study (Mark Welch et al., 2024, Figure 13) which documented <u>increased</u> transcripts of ribosomal proteins in undifferentiated HL-60/sh1 cells, in contrast to <u>decreased</u> transcripts in undifferentiated HL-60/S4 cells.

**Table 7.** GSEA results for the five different gene sets: KEGG; GOCC; ALL; ribosomal protein only (rp); non-ribosomal protein (nrp).

| CELLS | NES | Nominal p |
| --- | --- | --- |
| S4 RA/0 | -2.16 | 0.0 |
| sh1 RA/0 | 2.17 | 0.0 |
| gfp RA/0 | 1.72 | 0.0 |

GOCC\_RIBOSOME genes
| CELLS | NES | Nominal p |
| --- | --- | --- |
| S4 RA/0 | -2.26 | 0.0 |
| sh1 RA/0 | 1.13 | 0.17 |
| gfp RA/0 | -1.37 | 0.0 |

GO:0005840\_ALL genes
| CELLS | NES | Nominal p |
| --- | --- | --- |
| S4 RA/0 | -2.14 | 0.0 |
| sh1 RA/0 | 1.18 | 0.10 |
| gfp RA/0 | -1.33 | 0.01 |

**GO:0005840\_ ribosomal protein only****163 genes**
| CELLS | NES | Nominal p |
| --- | --- | --- |
| S4 RA/0 | -2.48 | 0.0 |
| sh1 RA/0 | 1.41 | 0.003 |
| gfp RA/0 | -1.24 | 0.02 |

| CELLS | NES | Nominal p |
| --- | --- | --- |
| S4 RA/0 | -1.04 | 0.37 |
| sh1 RA/0 | -1.40 | 0.02 |
| gfp RA/0 | -1.40 | 0.02 |

Despite the general agreement concerning the elevation of ribosomal structural protein transcripts (and presumably, ribosomal structural proteins), there remains a question whether there is an elevation of “completed” ribosomes. This predicament arises from GSEA analysis of ribosome (structural) RNA transcription. Table 8 demonstrates that in two analyses (GOBP_RRNA_TRANSCRIPTION and GOBP_NUCLEOLAR_LARGE_RRNA-_TRANSCRIPTION_BY_RNA_POLYMERASE_I) there are significant <u>decreases</u> of ribosome RNA transcription during RA-induced granulocyte differentiation of HL-60/S4, HL-60/sh1 and HL-60/gfp cells. Table S5 presents the GSEA Leading Edges of the downregulated ribosome RNA synthesis genes in the three differentiated cell lines. This predicament introduces the question: How are the levels of ribosomal proteins and RNAs coordinated? With regards to the synthesis of ribosomal proteins only, there are multiple genes with multiple regulators generating a “mind-boggling” complexity (Petibon et al., 2021). Coordination of ribosomal protein and RNA cellular levels seems to be controlled by a single gene (RPH1) in yeast (Shu et al., 2020); but similar control has not been described in human cells. Unfortunately, we are forced to speculate (without clear evidence) that the nucleoli of the three granulocytic HL-60 cell lines can synthesize enough ribosomal RNAs to generate functional ribosomes until their death by apoptosis.

**Table 8.**
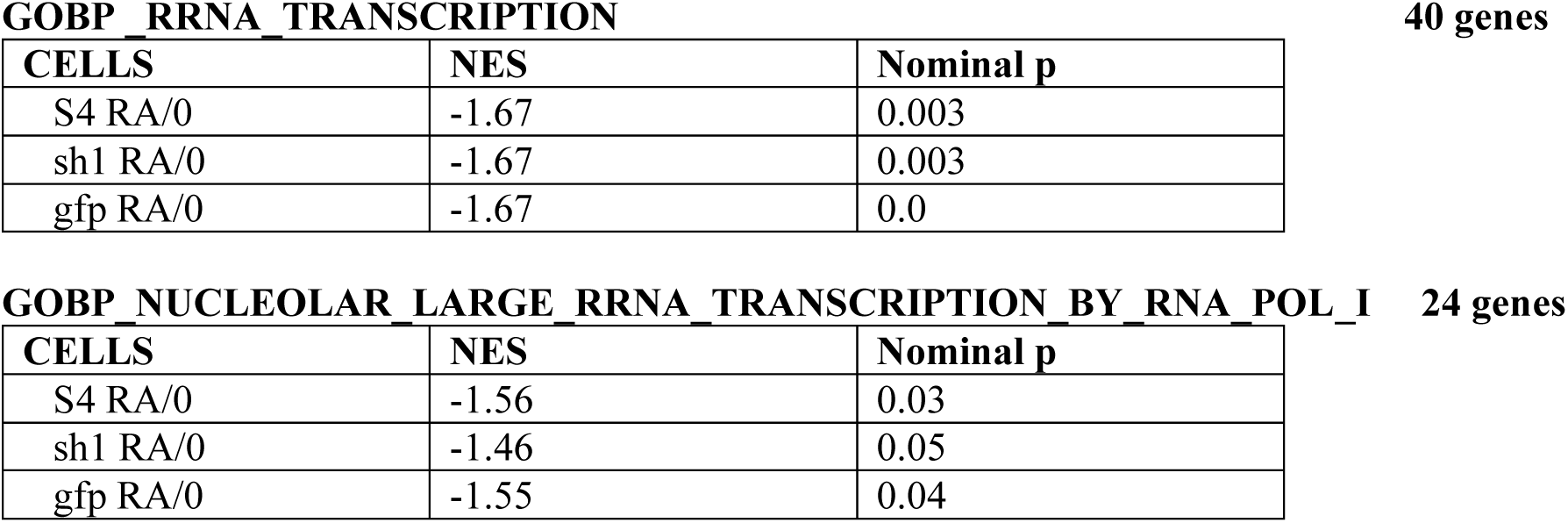
GSEA results for ribosome RNA transcription during RA-induced granulocyte differentiation of HL-60/S4, HL-60/sh1 and HL-60/gfp cells.

### Part One Summary

The effects of LBR knockdown upon granulocyte differentiation, comparing the transcriptomes (RA/0) of HL-60/S4, HL-60/sh1 and HL-60/gfp cell lines, revealed only minor phenotypic differences in predicted granulocyte behavior. Among the more significant consequences of LBR knockdown are the increased loss of heterochromatin and the decreased histone methyltransferase activity in RA-differentiated HL-60/sh1 cells. In addition, ribosome structural protein transcripts also appear to be increased during granulocyte differentiation, probably sufficient in amount for the limited granulocyte life-span.

### Part Two

#### Undifferentiated HL-60/sh1 cells are enriched in ribosome biosynthesis transcripts compared to undifferentiated HL-60/S4 cells

As mentioned earlier in this article, undifferentiated HL-60/S4 and HL-60/sh1 grow very robustly in vitro; whereas, granulocytic versions of the three cell lines HL-60/S4, HL-60/sh1 and HL-60/gfp exhibit enrichment of apoptosis genes (Figure 5). As a consequence, in order to investigate the role of LBR in ribosome biosynthesis of active cycling cells, we must compare undifferentiated HL-60/sh1 with undifferentiated HL-60/S4 cells (i.e., sh1 0 versus S4 0). Employing GOBP_RIBOSOME_ASSEMBLY and GOBP_RIBOSOME_BIOGENESIS (Figure 12) clearly indicates that LBR knockdown in undifferentiated HL-60/sh1 results in <u>enrichment</u> of gene transcripts supporting the considerable synthesis of the ribosomes.

**Figure 12.**
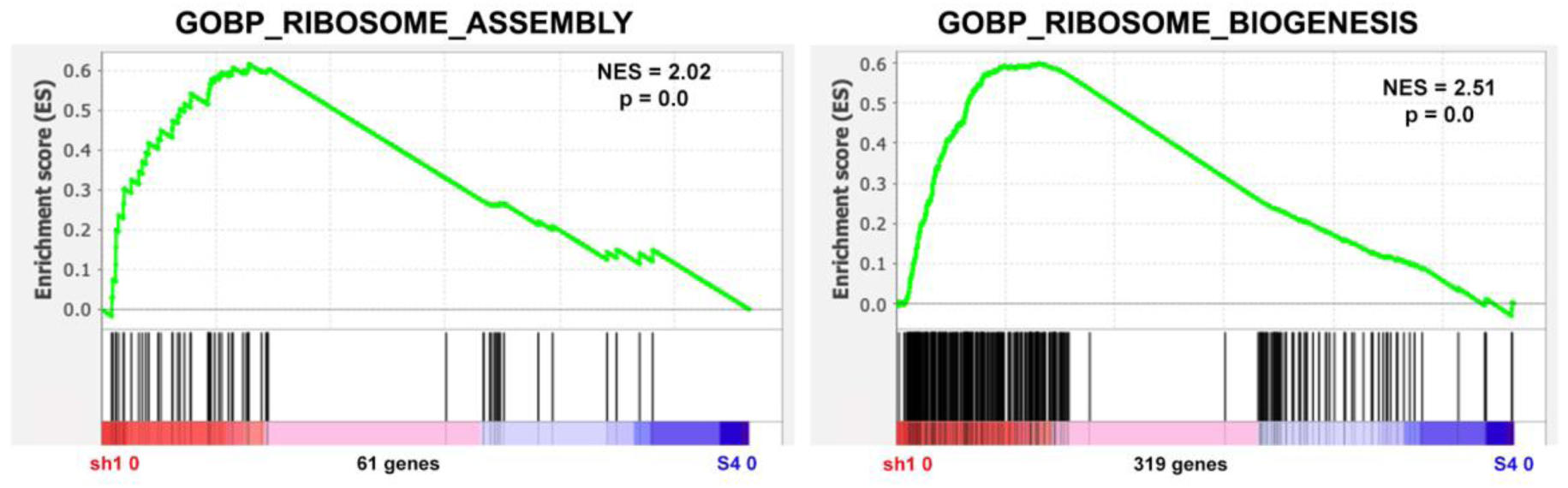
GSEA enrichment plots show enrichment of nucleolar ribosome assembly and biosynthesis transcripts in the sh1 0 cells compared to the S4 0 cells.

In **Part One** of this article, we discussed the problem of rRNA (Ribosome RNA) synthesis in the differentiated granulocytes (Table 8). This does not appear to be a problem, when we compare cycling HL-60/sh1 0 versus cycling HL-60/S4 (Figure 13 and Table 9). There is also significant enrichment in GOBP_RIBOSOMAL_SUBUNIT_EXPORT_FROM_NUCLEUS (Table 9, GSEA enrichment plot is not shown).

**Figure 13.**
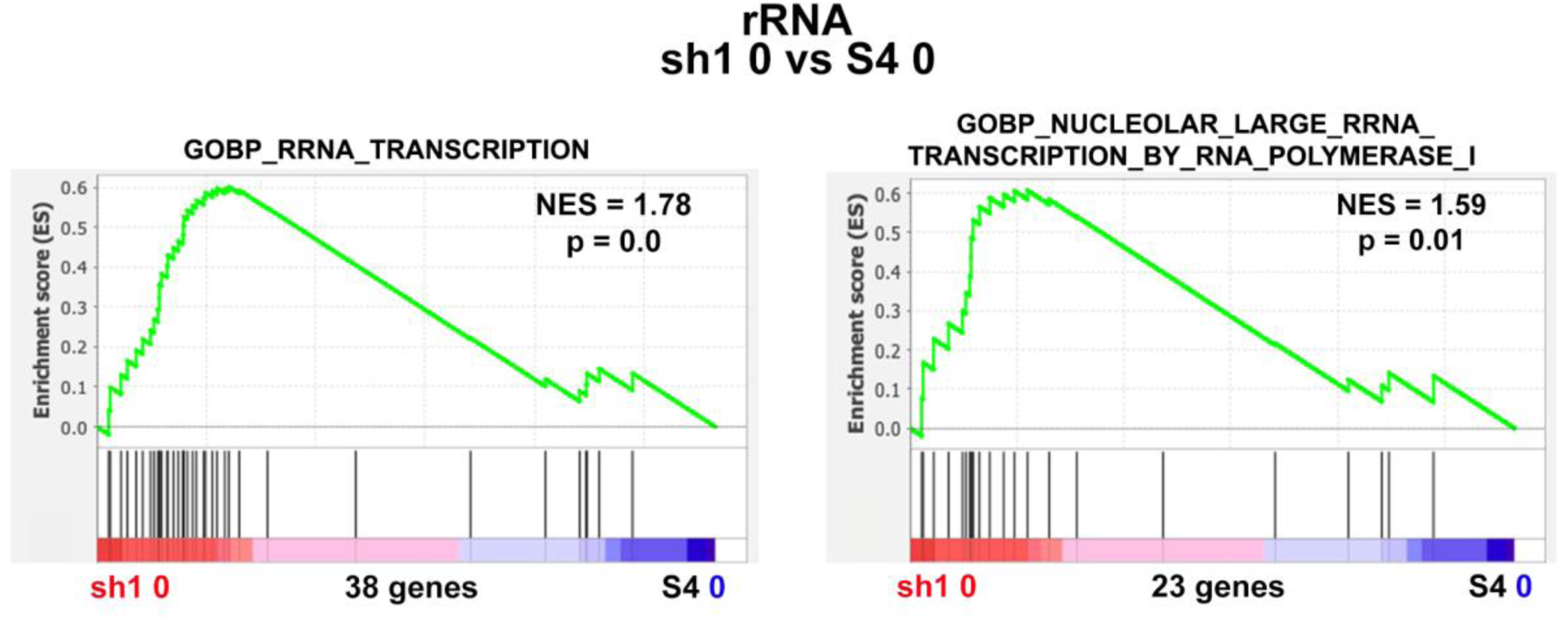
GSEA enrichment plots show enrichment of nucleolar large ribosome RNA biosynthesis transcripts in the sh1 0 cells compared to the S4 0 cells.

**Table 9.** GSEA results for ribosome biosynthesis, including ribosome RNA transcription, comparing HL-60/sh1 0 cells to HL-60/S4 0 cells.

| GSEA Gene Set | NES | Nominal<br>p |
| --- | --- | --- |
| GOBP_RIBOSOME_ASSEMBLY | 2.02 | 0.0 |
| GOBP_RIBOSOME_BIOGENESIS | 2.51 | 0.0 |
| GOBP_RRNA_TRANSCRIPTION | 1.78 | 0.0 |
| GOBP_NUCLEOLAR_LARGE_RRNA_TRANSCRIPTION_BY_RNA_POLYMERASE_I | 1.59 | 0.01 |
| GOBP_RIBOSOMAL_SUBUNIT_EXPORT_FROM_NUCLEUS | 1.49 | 0.05 |

#### Spatial organization of the nucleolus is regarded as critical to ribosome biosynthesis

Active cycling cells exhibit prominent nucleoli that can reform after each mitosis. The nucleolar substructure (including the Fibrillar Center, the Dense Fibrillar Component and the Granular Component) have been extensively studied (Ballmer et al., 2023; Panov et al., 2021; Potapova and Gerton, 2019). The nucleolus is regarded as a prime example of a non-membranous liquid-liquid phase separated (LLPS) cellular structure (Lafontaine et al., 2021; Yoneda et al., 2021). Ribosome protein transcripts are translated in the cytoplasm and the proteins transported into the nucleolus, where ribosomal RNA is synthesized, together forming ribosomal subunits which then are transported back to the cytoplasm for final ribosome assembly (Table 9). This complex sequence of events requires proper nucleolar spatial organization, which appears to be more enriched in sh1 0 cells compared to the S4 0 cells (Figure 14).

**Figure 14.**
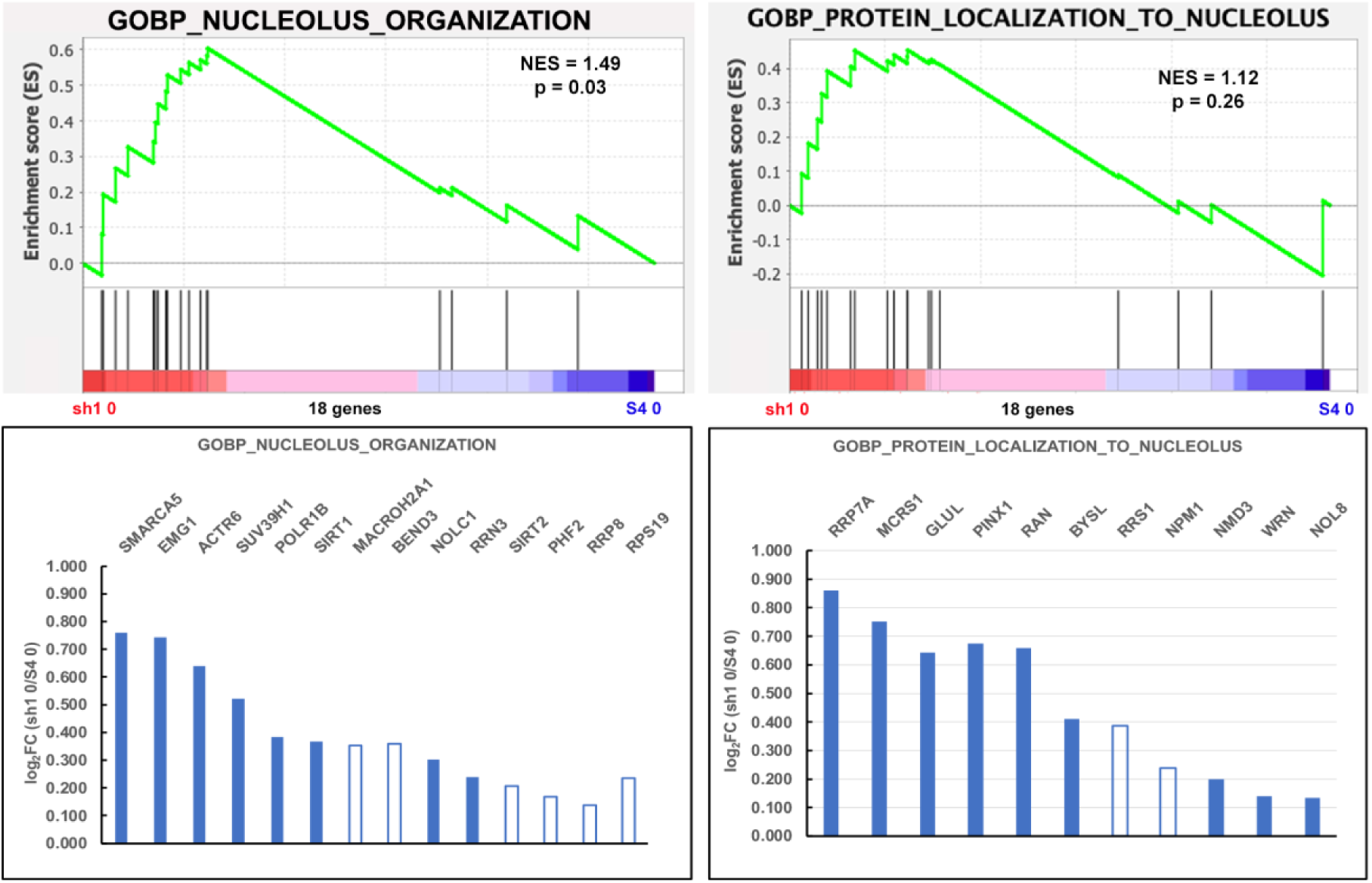
Expression of Nucleolus Genes. Top: GSEA enrichment plots of MSigDB GOBP_NUCLEOLUS_ORGANIZATION and GOBP_PROTEIN_LOCALIZATION-_TO_NUCLEOLUS gene sets showing increased enrichment in the sh1 0 cells compared to the S4 0 cells. Bottom: bar graphs of the DGE (Log_2_ FC) of the relevant Leading Edges. It is of interest to point out a few genes with known functions (www.genecards.org): POLR1B, RNA polymerase I subunit B; ACTR6, regulation of ribosomal DNA (rDNA) transcription; SMARCA5, chromatin remodeling to facilitate transcription. Closed bars: PPDE>0.95 (significant data). Open bars: PPDE<0.95 (not significant). HGNC Gene codes are displayed above the relative transcript level bars.

#### The texture of interphase chromatin, comparing undifferentiated HL-60/S4 to undifferentiated HL-60/sh1 cells, exhibits structural differences likely resulting from LBR knockdown

Employing DAPI to visualize DNA (as chromatin) density distribution in HCHO-fixed interphase nuclei reveals fluctuations in stain intensity, where the strongest staining (e.g., adjacent to the nuclear envelope and at the surface of nucleoli) can be regarded as heterochromatin. The current view of the formation of heterochromatin is that nucleosomal chromatin fibers are compacted into LLPS “condensates” by a combination of specific and unstructured interactions (e.g., HP1 proteins, DNA and histone epigenetic modifications) (Gibson et al., 2019; Narlikar, 2020; Olins and Olins, 2018; Zhang et al., 2022). A notable structural difference is observed in the perinucleolar chromatin (PNC) surrounding the nucleoli of undifferentiated HL-60/sh1 cells. In this cell state, the PNC often exhibits “bright” DAPI stained condensates and a thickened PNC layer adjacent to the round nucleolar “droplet” (Figure 15). In general, the interphase chromatin texture appears “smoother” in the HL-60/S4 cells, than in the HL-60/sh1 cells.

**Figure 15.**
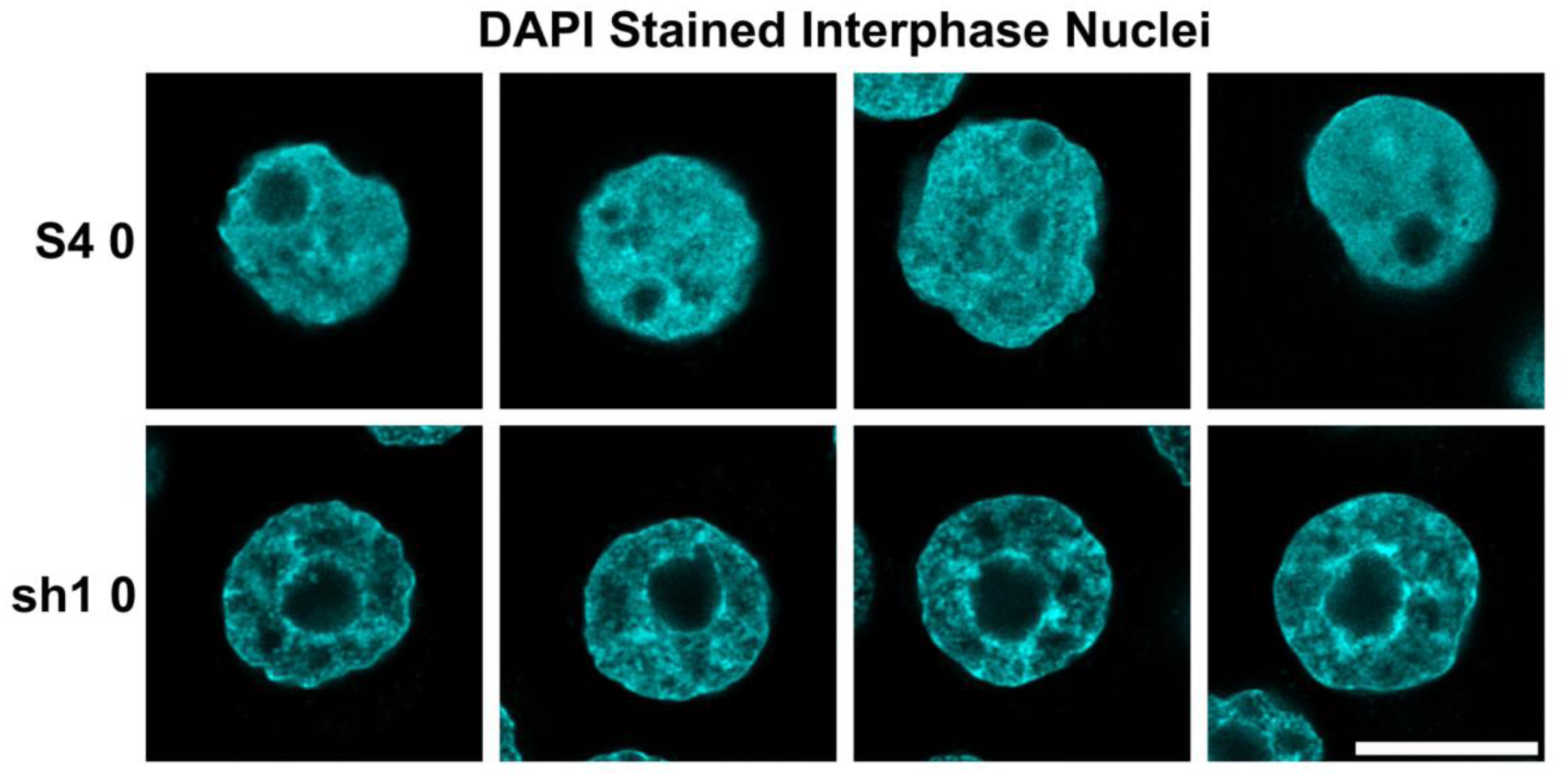
DAPI staining of undifferentiated interphase HL-60/S4 and HL-60/sh1 cells illustrating the “smooth” texture of the S4 0 interphase chromatin, compared to the “coarse” texture of sh1 0 interphase chromatin “condensates” around the nucleoli. Magnification bar: 10 μm.

As described in our earlier article on the effects of LBR knockdown on HL-60 nuclear architecture (Mark Welch et al., 2024), many sh1 0 cells display only one nucleolus, centrally located within the nucleus; whereas, S4 0 cells often possess multiple nucleoli, frequently in close proximity to the nuclear envelope. Figure 16 presents a quantitative assessment of this observation.

**Figure 16.**
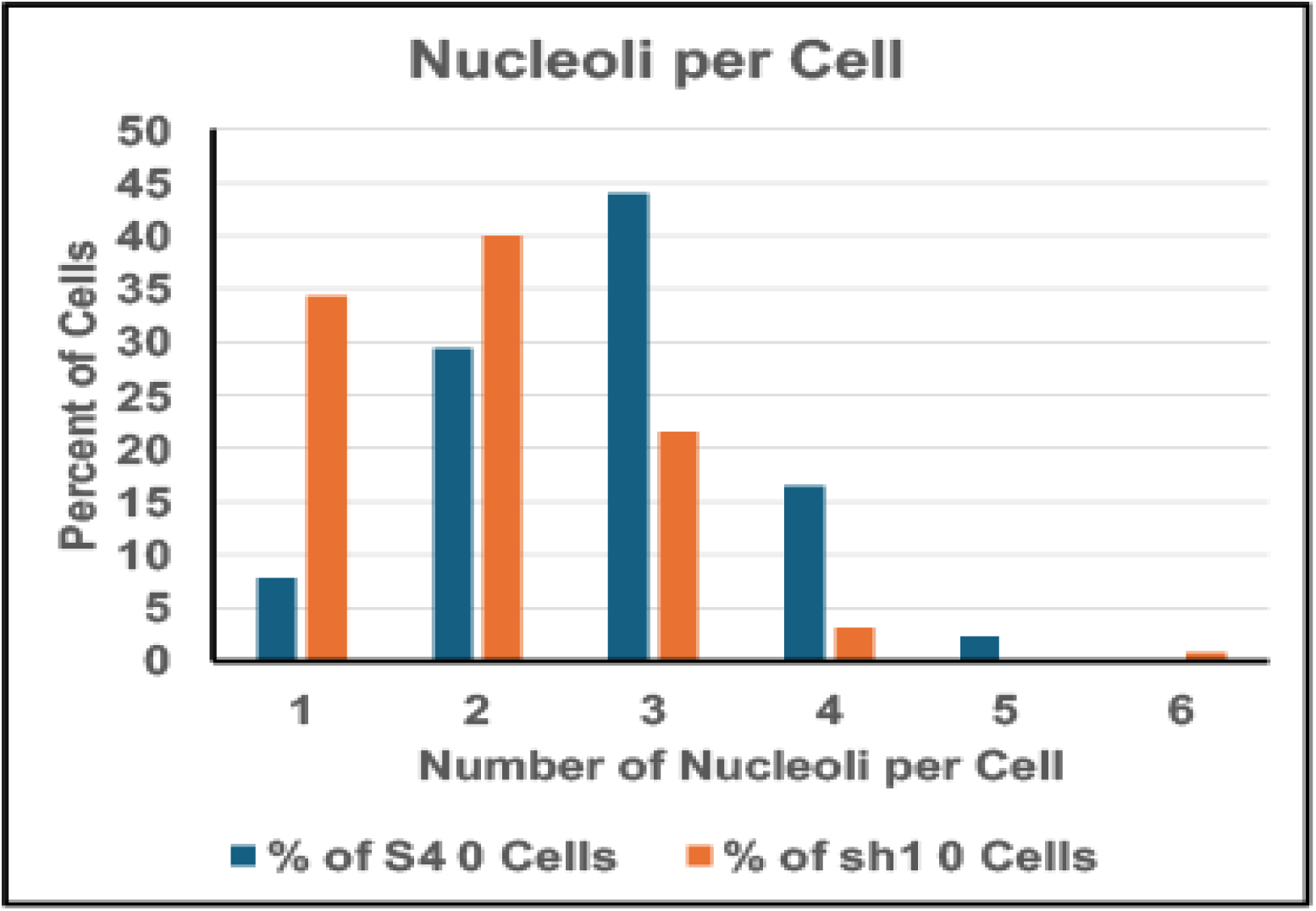
The percentage of cells with one-or-more nucleoli per cell. The number of cells counted (N) equals: S4 0 (218) and sh1 0 (125).

Another observation, made during the flourescent microscopic comparison of DAPI stained cycling HL-60/S4 0 cells versus cycling HL-60/sh1 0 cells is the difference in “roundness” of the interphase nuclei (Figures 15 and 17). It appears that a higher % of sh1 0 cells exhibit a round nucleus (∼68%; N=199), than estimated in S4 0 nuclei (∼22%; N=157). Most of the HL-60/S4 nuclei exhibit irregular shapes. Exactly how this difference in interphase nuclear shape results from the presence-or-absence of LBR remains to be illucidated. We speculate that in the absence of LBR, the nuclear envelope may reveal an intrinsic uniform curvature. This intrinsic curvature may arise from a uniform distribution of specific lipids and/or a uniform distribution of nuclear pores (Peeters et al., 2022). In HL-60/S4 cells, the presence of LBR may result in attachment of diverse regions of heterochromatin to the nuclear envelope, resulting in complicated nuclear shapes.

**Figure 17.**
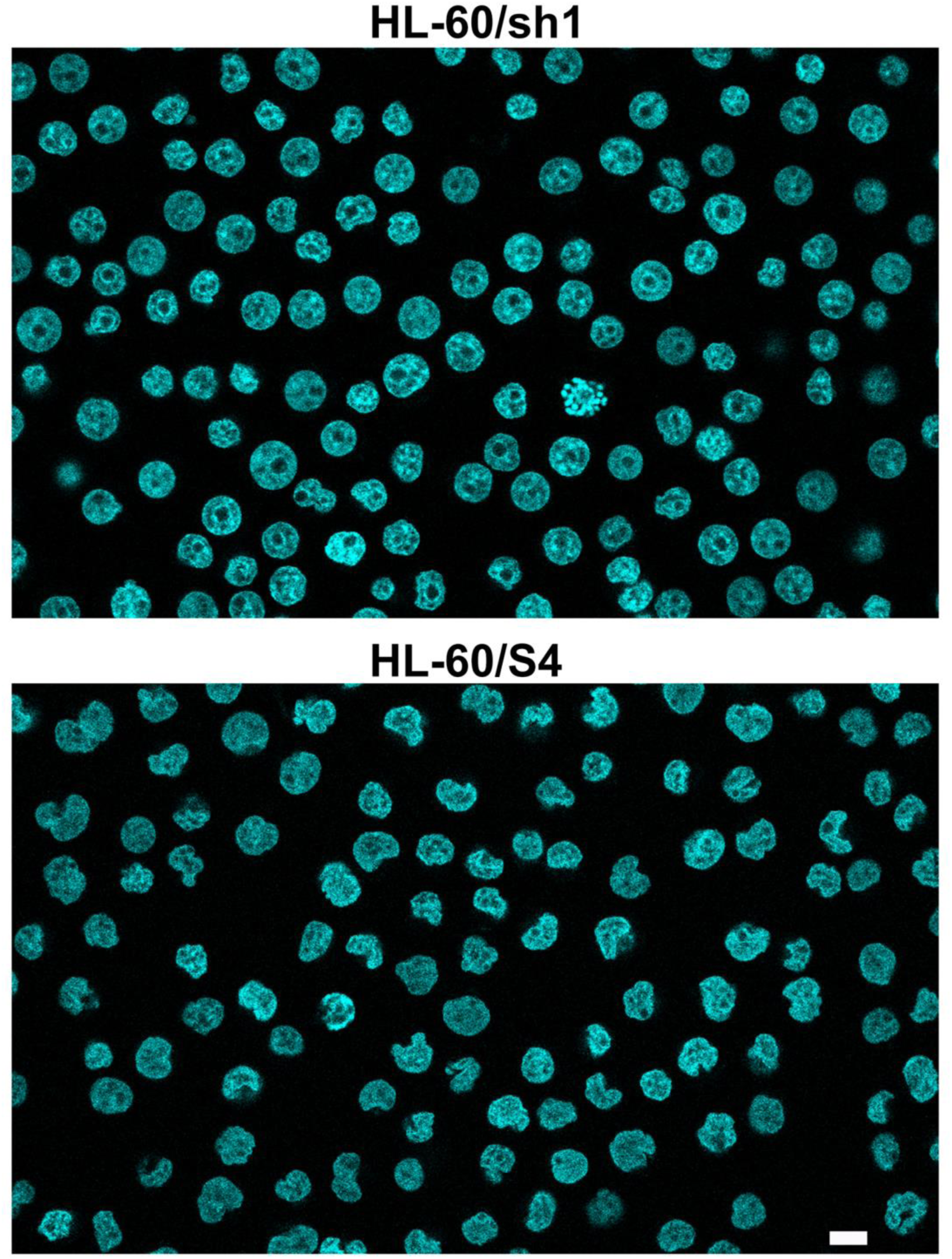
Comparison of the nuclear shapes of DAPI stained cycling HL-60/sh1 0 (Top Panel) versus cycling HL-60/S4 0 (Bottom Panel). Magnification bar: 10 μm.

The prevalent nucleolar “central position” and the increased “roundness” of undifferentiated HL-60/sh1 nuclei could also be influenced by: 1) the LINC (“Linker of Nucleoskeleton and Cytoskeleton”) Complex; 2) the LEM (LAP2-emerin-MAN1) domain proteins; 3) the PLEC-VIM (Plectin-Vimentin) intermediate filament connections to the interphase nuclear envelope. These interactions with the nuclear envelope are described in excellent reviews (Barton et al., 2015; Djabali, 1999; King, 2023). In order to formulate a hypothesis on the contributions of these structural systems, it is necessary to see how they differ in gene expression of key genes within undifferentiated HL-60/S4, HL-60/sh1 and HL-60/gfp cell lines that differ in expression of LBR (Figure 18). Of particular interest is the significant downregulation within undifferentiated HL-60/sh1 cells of the LINC participant SYNE1 (Nesprin1). The relative (sh1/S4) gene expression of SYNE1: Log_2_FC is -3.00 or an 8-fold reduction in SYNE1 transcripts. This significant SYNE1 downregulation might “weaken” or “break” the LINC complex that spans: Actin Cytoskeleton-Nesprin-Outer Nuclear Membrane-SUN-Inner Nuclear Membrane-Lamins A/C and B (King, 2023). A similar observation has been made with breast cancer cells, where SUN1 was knocked down (Matsumoto et al., 2016). In the situation of HL-60/sh1 cells, it appears that LBR knockdown is important for the significant reduction of SYNE1. We can not rule out contributions from LEM-D and IF gene expression changes in this acquired phenotypic state.

**Figure 18.**
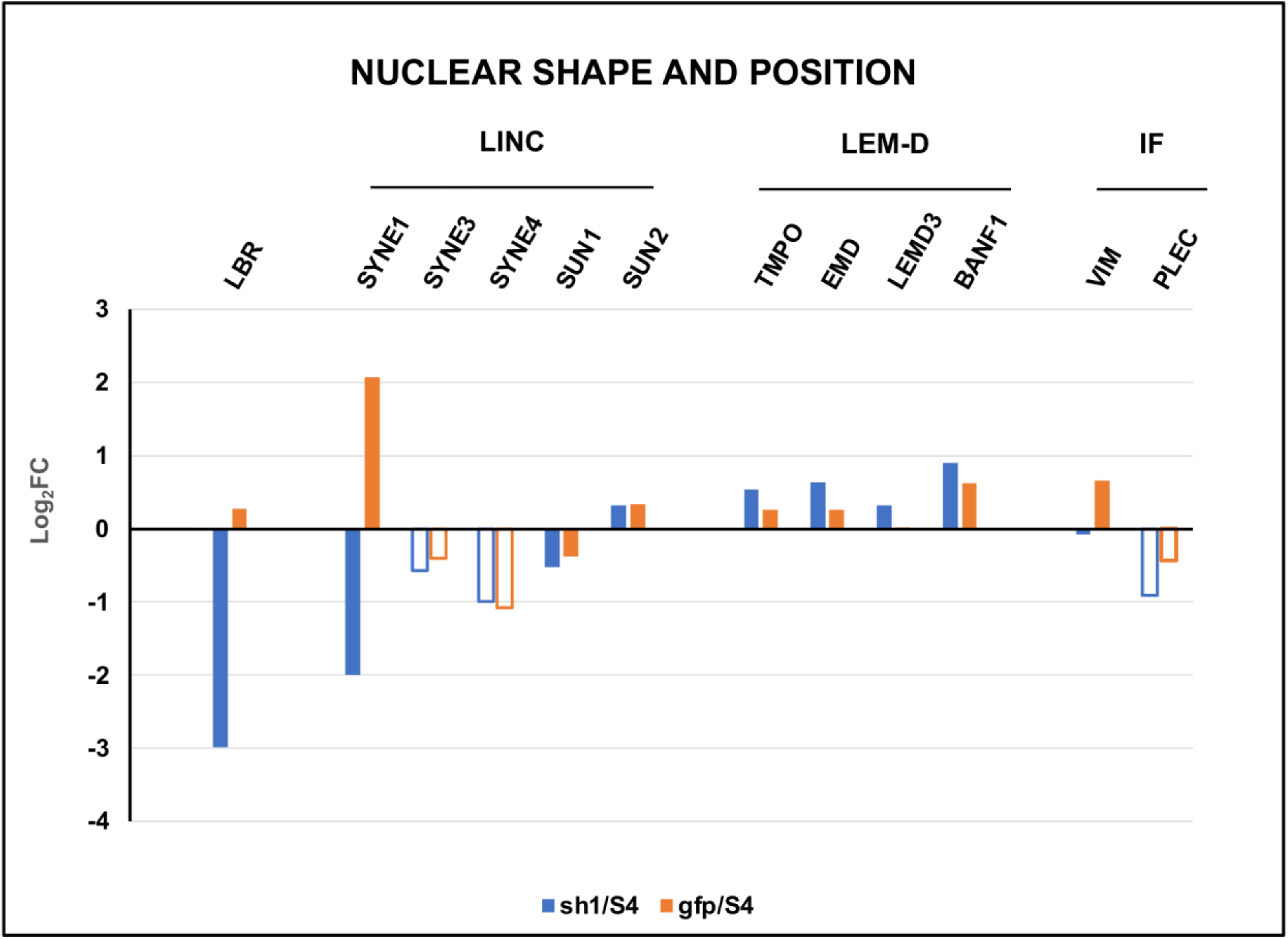
Differential gene expression of some key genes likely involved in nuclear shape determination and nuclear position in the cell. Gene Categories: LINC, Linker of Nucleoskeleton and Cytoskeleton Complex; LEM-D, LEM domain proteins; IF, intermediate filament proteins. LBR is included for comparison.

#### Heterochromatin and Nucleosome Formation appear to be increased in undifferentiated HL-60/sh1 cells compared to undifferentiated HL-60/S4 cells

In our earlier article (Olins et al., 2024), which explored the phenotypes of senescent HL-60/sh1 macrophages, we noted a GSEA <u>reduction</u> of heterochromatin and nucleosome transcripts in these TPA treated HL-60/S4 cells compared to untreated HL-60/S4 cells. By contrast, in the present study, comparing undifferentiated HL-60/sh1 to undifferentiated HL-60/S4, we observe significant GSEA <u>enrichment</u> of heterochromatin and nucleosome transcripts in the HL-60/sh1 cells (Figures 19, 20 and Table 10).

**Figure 19.**
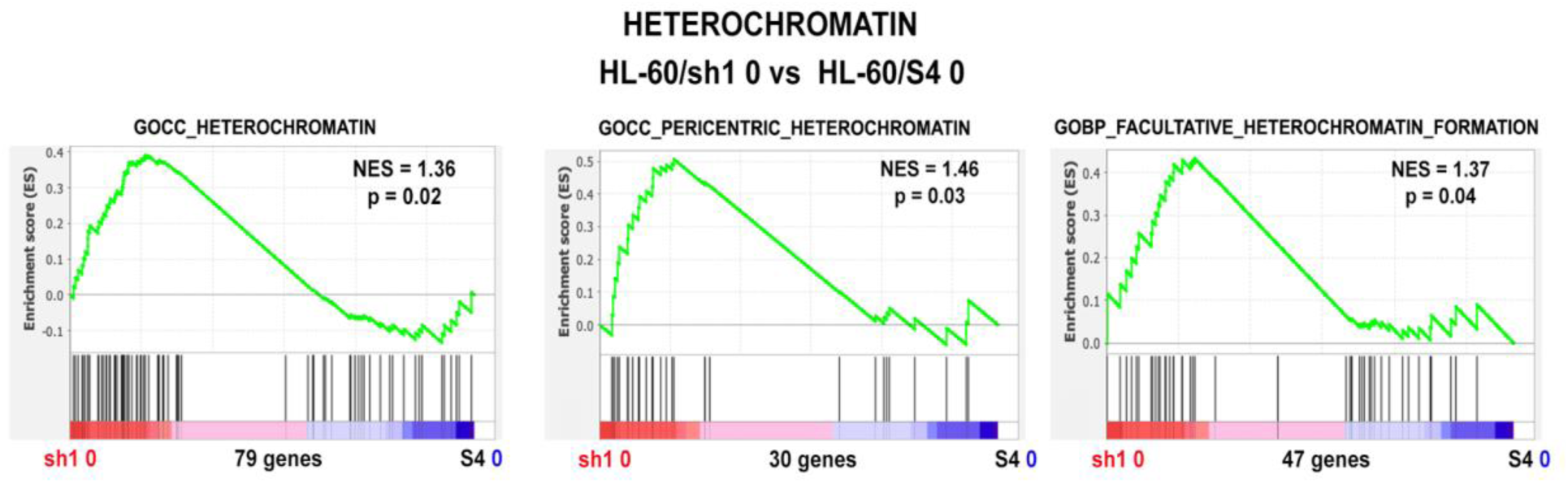
GSEA enrichment plots of three MSigDB gene sets demonstrating that sh1 0 cells exhibit a greater enrichment in heterochromatin-related transcripts, than S4 0 cells. Four genes are shared in the Leading Edges: SMARCA5, HELLS, EZH2 and MACROH2A1.

**Figure 20.**
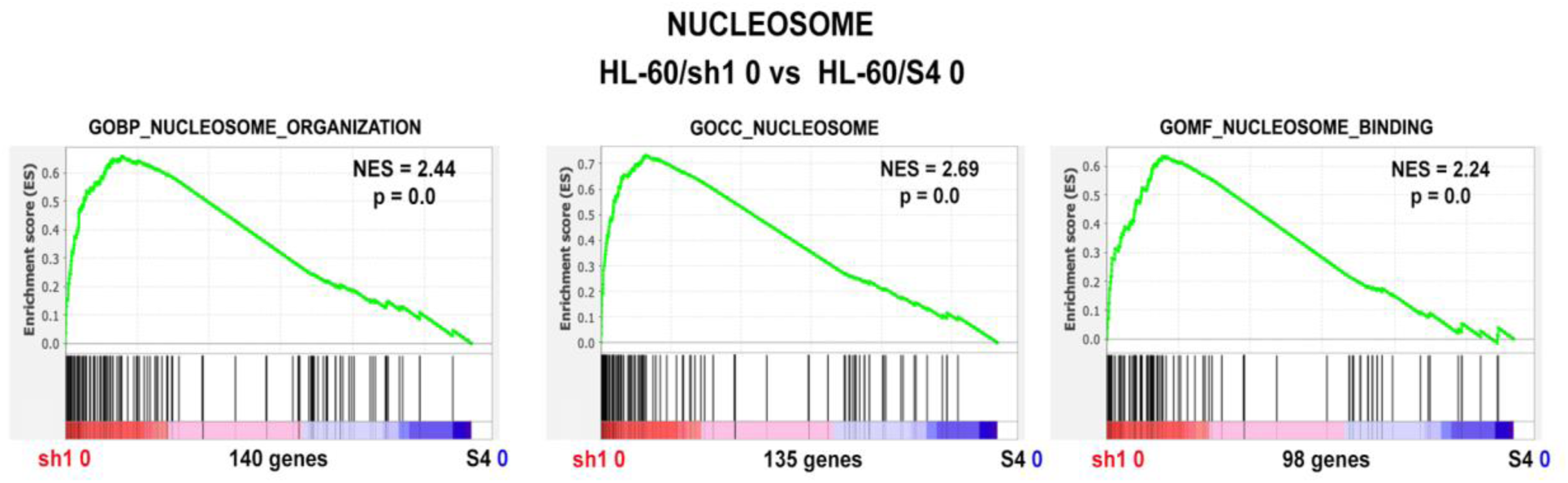
GSEA enrichment plots of three MSigDB gene sets involving the nucleosome, demonstrating that sh1 0 cells exhibit a greater enrichment in nucleosome-related transcripts than S4 0 cells. This enrichment is demonstrated in three different GSEA gene sets. Four genes are shared in the Leading Edges: H1.5, H1.4, H3-3B and H2AX.

**Table 10.** GSEA results of additional MSigDB gene sets involving heterochromatin and the nucleosome.

| GSEA Gene Set | NES | Nominal p |
| --- | --- | --- |
| GOCC_HETEROCHROMATIN | 1.36 | 0.02 |
| GOCC_PERICENTRIC_HETEROCHROMATIN | 1.46 | 0.03 |
| GOBP_FACULTATIVE_HETEROCHROMATIN_FORMATION | 1.37 | 0.04 |
| GOBP_NUCLEOSOME_ORGANIZATION | 2.44 | 0.0 |
| GOCC_NUCLEOSOME | 2.69 | 0.0 |
| GOMF_NUCLEOSOME_BINDING | 2.24 | 0.0 |

### Part Two Summary

Comparing the transcriptomes and DAPI staining of undifferentiated HL-60/sh1 and HL-60/S4 cell lines (i.e., sh1 0 versus S4 0), we observed a number of surprises paralleling the LBR knockdown. In contrast to a <u>deficit</u> of ribosomal RNA synthesized during granulocyte differentiation, we noted a significant <u>increase</u> in ribosomal protein and ribosomal RNA synthesis in sh1 0 cells, compared to the S4 0 cell line. Also, we noted a significant increase in heterochromatin and nucleosome formation in sh1 0 compared to S4 0 cells. This latter observation may be related to the dramatic morphological changes in DAPI staining of sh1 0 nuclei, which appeared to show chromatin condensates surrounding nucleoli. Furthermore, the sh1 0 nuclei appeared to be “rounder” than the more “contorted” S4 0 nuclei. We present transcriptome evidence that the sh1 0 cells may have deficiencies in the LINC complex and intermediate filament connections with the nuclear envelope.

## Discussion

The purpose of the present and predecessor articles (Mark Welch et al., 2024; Olins et al., 2010a) is to understand the cellular functions of Lamin B Receptor (LBR) in the myeloid HL-60/S4 cell line by extensive analysis of an LBR knockdown (HL-60/sh1) cell line. Prior knowledge on the structure and functions of LBR has been summarized (Giannios et al., 2017; Nikolakaki et al., 2017; Olins et al., 2010b). It is known that in HL-60/S4 cells, LBR is responsible for nuclear lobulation in RA-differentiated granulocytes (Olins et al., 1998; Olins et al., 2001). It is generally accepted that LBR connects heterochromatin to the interphase nuclear envelope and also functions as an essential enzyme in cholesterol biosynthesis (Giannios et al., 2017; Nikolakaki et al., 2017; Olins et al., 2010b).

In the preceding study (Mark Welch et al., 2024) and in the first part of the present study, we have focused on the consequences of LBR knockdown upon aspects of RA-induced granulocyte differentiation. Most of the analyses were of relative transcript levels in differentiated versus undifferentiated (RA/0) cells, comparing HL-60/sh1 and HL-60/S4 cells. Examining the Log_2_FC (Fold Changes) of RA/0, we were able to demonstrate that many of the Golgi Apparatus protein transcripts were downregulated and many of the ribosome protein transcripts were upregulated during RA-induced differentiation of HL-60/sh1, but not HL-60/S4 cells. Examination of the Golgi proteins led to identifying downregulation of TRIP11, a candidate gene for a human chondrodysplasia, which is a phenocopy for Greenberg Dysplasia, a chondrodysplasia arising from LBR mutations (Wehrle et al., 2018). We (Mark Welch et al., 2024) also suggested that the nucleolar changes in HL-60/sh1 cells may be related to the increase of ribosome protein transcripts. The deficiency of LBR may allow nucleolar (droplet) fusion leading to more efficient rRNA synthesis in a Liquid-Liquid Phase Separated (LLPS) centrally-located nucleolus.

The present study employed GSEA (Gene Set Enrichment Analysis), previously utilized to study the transcriptome of TPA-differentiated HL-60/S4 macrophages (Olins et al., 2024), to examine the phenotypes of RA-differentiated granulocytes. Among the more surprising of conclusions about these granulocytes, the comparative transcriptomes imply that the three cell lines (HL-60/S4, HL-60/sh1 and HL-60/gfp), possess similar phenotypes, including enriched pro-apoptotic transcriptomes. Also of interest, comparing the three cell lines with three separate GSEA heterochromatin gene sets, we demonstrated that the RA-<u>differentiated</u> granulocytes all exhibited <u>decreased</u> heterochromatin, with HL-60/sh1 cells showing the greatest decrease. Comparing the Leading Edges for the three separate gene sets allowed identification of 8 genes, whose decreased transcript levels may be responsible for the greater loss of heterochromatin in HL-60/sh1 granulocytes. By contrast, in the second part of this article, comparing the three <u>undifferentiated</u> cell lines using the same GSEA heterochromatin gene sets, we observed <u>increased</u> heterochromatin, with HL-60/sh1 cells showing the greatest increase.

Based upon these recent studies of the three derived HL-60 cell lines, it is abundantly clear that asking the question, “What is the function of LBR?”, is <u>not</u> a simple question. Knockdown of LBR (e.g., HL-60/sh1 cells) displays different phenotypic consequences in differentiated versus undifferentiated cells. At the minimum, LBR connects heterochromatin to the interphase nuclear envelope. Furthermore, LBR appears to be important in associating nucleoli with the nuclear envelope and, possibly, interfering with nucleolar fusion. Although LBR is not regarded as a specific transcription repressor, it’s association with CBX1, CBX3 and CBX5 heterochromatin proteins suggests that it might affect many diverse genes, as well as influencing chromatin higher-order structure. In addition, the role of LBR in sterol biosynthesis suggests that sterol levels and locations might affect cell membranes, including the Golgi apparatus. It is evident that understanding the multiple functions of LBR will remain a field for continuing probing research.

## Acknowledgements

ALO and DEO express our gratitude to the MaineHealth Institute for Research for allowing us to join the research group of Dr. Igor Prudovsky. We thank Dr. Prudovsky for his generous hospitality in his laboratory.

## References

1. Ballmer, D., M. Tardat, R. Ortiz, A. Graff-Meyer, E.A. Ozonov, C. Genoud, A.H. Peters, and G. Fanourgakis. 2023. HP1 proteins regulate nucleolar structure and function by secluding pericentromeric constitutive heterochromatin. Nucleic Acids Res. 51:117–143.

2. Barton, L.J., A.A. Soshnev, and P.K. Geyer. 2015. Networking in the nucleus: a spotlight on LEM-domain proteins. Curr Opin Cell Biol. 34:1–8.

3. Djabali, K. 1999. Cytoskeletal proteins connecting intermediate filaments to cytoplasmic and nuclear periphery. Histol Histopathol. 14:501–509.

4. Fukuda, K., T. Shimi, C. Shimura, T. Ono, T. Suzuki, K. Onoue, S. Okayama, H. Miura, I. Hiratani, K. Ikeda, Y. Okada, N. Dohmae, S. Yonemura, A. Inoue, H. Kimura, and Y. Shinkai. 2023. Epigenetic plasticity safeguards heterochromatin configuration in mammals. Nucleic Acids Res. 51:6190–6207.

5. Giannios, I., E. Chatzantonaki, and S. Georgatos. 2017. Dynamics and Structure-Function Relationships of the Lamin B Receptor (LBR). PLoS One. 12:e0169626.

6. Gibson, B.A., L.K. Doolittle, M.W.G. Schneider, L.E. Jensen, N. Gamarra, L. Henry, D.W. Gerlich, S. Redding, and M.K. Rosen. 2019. Organization of Chromatin by Intrinsic and Regulated Phase Separation. Cell. 179:470–484.e421.

7. Hoffmann, K., C.K. Dreger, A.L. Olins, D.E. Olins, L.D. Shultz, B. Lucke, H. Karl, R. Kaps, D. Müller, A. Vayá, J. Aznar, R.E. Ware, N. Sotelo Cruz, T.H. Lindner, H. Herrmann, A. Reis, and K. Sperling. 2002. Mutations in the gene encoding the lamin B receptor produce an altered nuclear morphology in granulocytes (Pelger-Huët anomaly). Nat Genet. 31:410–414.

8. King, M.C. 2023. Dynamic regulation of LINC complex composition and function across tissues and contexts. FEBS Lett. 597:2823–2832.

9. Lafontaine, D.L.J., J.A. Riback, R. Bascetin, and C.P. Brangwynne. 2021. The nucleolus as a multiphase liquid condensate. Nat Rev Mol Cell Biol. 22:165–182.

10. Leung, M.F., J.A. Sokoloski, and A.C. Sartorelli. 1992. Changes in microtubules, microtubule-associated proteins, and intermediate filaments during the differentiation of HL-60 leukemia cells. Cancer Res. 52:949–954.

11. Mark Welch, D.B., A. Jauch, J. Langowski, A.L. Olins, and D.E. Olins. 2017. Transcriptomes reflect the phenotypes of undifferentiated, granulocyte and macrophage forms of HL-60/S4 cells. Nucleus. 8:222–237.

12. Mark Welch, D.B., A.L. Olins, and D.E. Olins. 2024. The Effects of Lamin B Receptor knockdown on a Myeloid Leukemia Cell. bioRxiv:2024.2006.2019.598074.

13. Matsumoto, A., C. Sakamoto, H. Matsumori, J. Katahira, Y. Yasuda, K. Yoshidome, M. Tsujimoto, I.G. Goldberg, N. Matsuura, M. Nakao, N. Saitoh, and M. Hieda. 2016. Loss of the integral nuclear envelope protein SUN1 induces alteration of nucleoli. Nucleus. 7:68–83.

14. Narlikar, G.J. 2020. Phase-separation in chromatin organization. J Biosci. 45.

15. Nikolakaki, E., I. Mylonis, and T. Giannakouros. 2017. Lamin B Receptor: Interplay between Structure, Function and Localization. Cells. 6.

16. Olins, A.L., B. Buendia, H. Herrmann, P. Lichter, and D.E. Olins. 1998. Retinoic acid induction of nuclear envelope-limited chromatin sheets in HL-60. Exp Cell Res. 245:91–104.

17. Olins, A.L., A. Ernst, M. Zwerger, H. Herrmann, and D.E. Olins. 2010a. An in vitro model for Pelger-Huët anomaly: stable knockdown of lamin B receptor in HL-60 cells. Nucleus. 1:506–512.

18. Olins, A.L., H. Herrmann, P. Lichter, M. Kratzmeier, D. Doenecke, and D.E. Olins. 2001. Nuclear envelope and chromatin compositional differences comparing undifferentiated and retinoic acid- and phorbol ester-treated HL-60 cells. Exp Cell Res. 268:115–127.

19. Olins, A.L., T.V. Hoang, M. Zwerger, H. Herrmann, H. Zentgraf, A.A. Noegel, I. Karakesisoglou, D. Hodzic, and D.E. Olins. 2009. The LINC-less granulocyte nucleus. Eur J Cell Biol. 88:203–214.

20. Olins, A.L., D. Mark Welch, D. Saul, I. Prudovsky, and D.E. Olins. 2024. The Multifaceted Phenotype of Senescent HL-60/S4 Macrophages. bioRxiv:2024.2006.2015.598082.

21. Olins, A.L., G. Rhodes, D.B. Welch, M. Zwerger, and D.E. Olins. 2010b. Lamin B receptor: multi-tasking at the nuclear envelope. Nucleus. 1:53–70.

22. Olins, D.E., and A.L. Olins. 2018. Epichromatin and chromomeres: a ’fuzzy’ perspective. Open Biol. 8.

23. Padeken, J., S.P. Methot, and S.M. Gasser. 2022. Establishment of H3K9-methylated heterochromatin and its functions in tissue differentiation and maintenance. Nat Rev Mol Cell Biol. 23:623–640.

24. Panov, K.I., K. Hannan, R.D. Hannan, and N. Hein. 2021. The Ribosomal Gene Loci-The Power behind the Throne. Genes (Basel). 12.

25. Parry, A.J., M. Hoare, D. Bihary, R. Hänsel-Hertsch, S. Smith, K. Tomimatsu, E. Mannion, A. Smith, P. D’Santos, I.A. Russell, S. Balasubramanian, H. Kimura, S.A. Samarajiwa, and M. Narita. 2018. NOTCH-mediated non-cell autonomous regulation of chromatin structure during senescence. Nat Commun. 9:1840.

26. Peeters, B.W.A., A.C.A. Piët, and M. Fornerod. 2022. Generating Membrane Curvature at the Nuclear Pore: A Lipid Point of View. Cells. 11.

27. Petibon, C., M. Malik Ghulam, M. Catala, and S. Abou Elela. 2021. Regulation of ribosomal protein genes: An ordered anarchy. Wiley Interdiscip Rev RNA. 12:e1632.

28. Potapova, T.A., and J.L. Gerton. 2019. Ribosomal DNA and the nucleolus in the context of genome organization. Chromosome Res. 27:109–127.

29. Rocha, A., A. Dalgarno, and N. Neretti. 2022. The functional impact of nuclear reorganization in cellular senescence. Brief Funct Genomics. 21:24–34.

30. Saul, D., R.L. Kosinsky, E.J. Atkinson, M.L. Doolittle, X. Zhang, N.K. LeBrasseur, R.J. Pignolo, P.D. Robbins, L.J. Niedernhofer, Y. Ikeno, D. Jurk, J.F. Passos, L.J. Hickson, A. Xue, D.G. Monroe, T. Tchkonia, J.L. Kirkland, J.N. Farr, and S. Khosla. 2022. A new gene set identifies senescent cells and predicts senescence-associated pathways across tissues. Nat Commun. 13:4827.

31. Shu, W.J., R. Chen, Z.H. Yin, F. Li, H. Zhang, and H.N. Du. 2020. Rph1 coordinates transcription of ribosomal protein genes and ribosomal RNAs to control cell growth under nutrient stress conditions. Nucleic Acids Res. 48:8360–8373.

32. Sławińska, N., and R. Krupa. 2021. Molecular Aspects of Senescence and Organismal Ageing-DNA Damage Response, Telomeres, Inflammation and Chromatin. Int J Mol Sci. 22.

33. Swanson, E.C., L.M. Rapkin, D.P. Bazett-Jones, and J.B. Lawrence. 2015. Unfolding the story of chromatin organization in senescent cells. Nucleus. 6:254–260.

34. Turner, E.M., and C. Schlieker. 2016. Pelger-Huët anomaly and Greenberg skeletal dysplasia: LBR-associated diseases of cholesterol metabolism. Rare Dis. 4:e1241363.

35. Wang, Z., and B. Ren. 2024. Role of H3K4 monomethylation in gene regulation. Curr Opin Genet Dev. 84:102153.

36. Wehrle, A., T.M. Witkos, J.C. Schneider, A. Hoppmann, S. Behringer, A. Köttgen, M. Elting, J. Spranger, M. Lowe, and E. Lausch. 2018. A common pathomechanism in GMAP-210- and LBR-related diseases. JCI Insight. 3.

37. Yoneda, M., T. Nakagawa, N. Hattori, and T. Ito. 2021. The nucleolus from a liquid droplet perspective. J Biochem. 170:153–162.

38. Young, A.N., E. Perlas, N. Ruiz-Blanes, A. Hierholzer, N. Pomella, B. Martin-Martin, A. Liverziani, J.W. Jachowicz, T. Giannakouros, and A. Cerase. 2021. Deletion of LBR N-terminal domains recapitulates Pelger-Huet anomaly phenotypes in mouse without disrupting X chromosome inactivation. Commun Biol. 4:478.

39. Zhang, H., H. Romero, A. Schmidt, K. Gagova, W. Qin, B. Bertulat, A. Lehmkuhl, M. Milden, M. Eck, T. Meckel, H. Leonhardt, and M.C. Cardoso. 2022. MeCP2-induced heterochromatin organization is driven by oligomerization-based liquid-liquid phase separation and restricted by DNA methylation. Nucleus. 13:1–34.

40. Zhang, X., X. Liu, Z. Du, L. Wei, H. Fang, Q. Dong, J. Niu, Y. Li, J. Gao, M.Q. Zhang, W. Xie, and X. Wang. 2021. The loss of heterochromatin is associated with multiscale three-dimensional genome reorganization and aberrant transcription during cellular senescence. Genome Res. 31:1121–1135.

